# Biomolecular condensates control and are defined by RNA-RNA interactions that arise in viral replication

**DOI:** 10.1101/2024.12.23.630161

**Authors:** Dilimulati Aierken, Vita Zhang, Rachel Sealfon, John C. Marecki, Kevin D. Raney, Amy S. Gladfelter, Jerelle A. Joseph, Christine A. Roden

## Abstract

Cells must limit RNA–RNA interactions to avoid irreversible RNA entanglement. Cells may prevent deleterious RNA-RNA interactions by genome organization to avoid complementarity however, RNA viruses generate long, perfectly complementary antisense RNA during replication. How do viral RNAs avoid irreversible entanglement? One possibility is RNA sequestration into biomolecular condensates. To test this, we reconstituted critical SARS-CoV-2 RNA–RNA interactions in Nucleocapsid condensates. We observed that RNAs with low propensity RNA–RNA interactions resulted in more round, liquid-like condensates while those with high sequence complementarity resulted in more heterogeneous networked morphology independent of RNA structure stability. Residue-resolution molecular simulations and direct sequencing-based detection of RNA–RNA interactions support that these properties arise from degree of trans RNA contacts. We propose that extensive RNA–RNA interactions in cell and viral replication are controlled via a combination of genome organization, timing, RNA sequence content, RNA production ratios, and emergent biomolecular condensate material properties.

**Graphical Abstract: SARS-CoV-2 replication cycle employs weak and strong RNA-RNA interactions:** 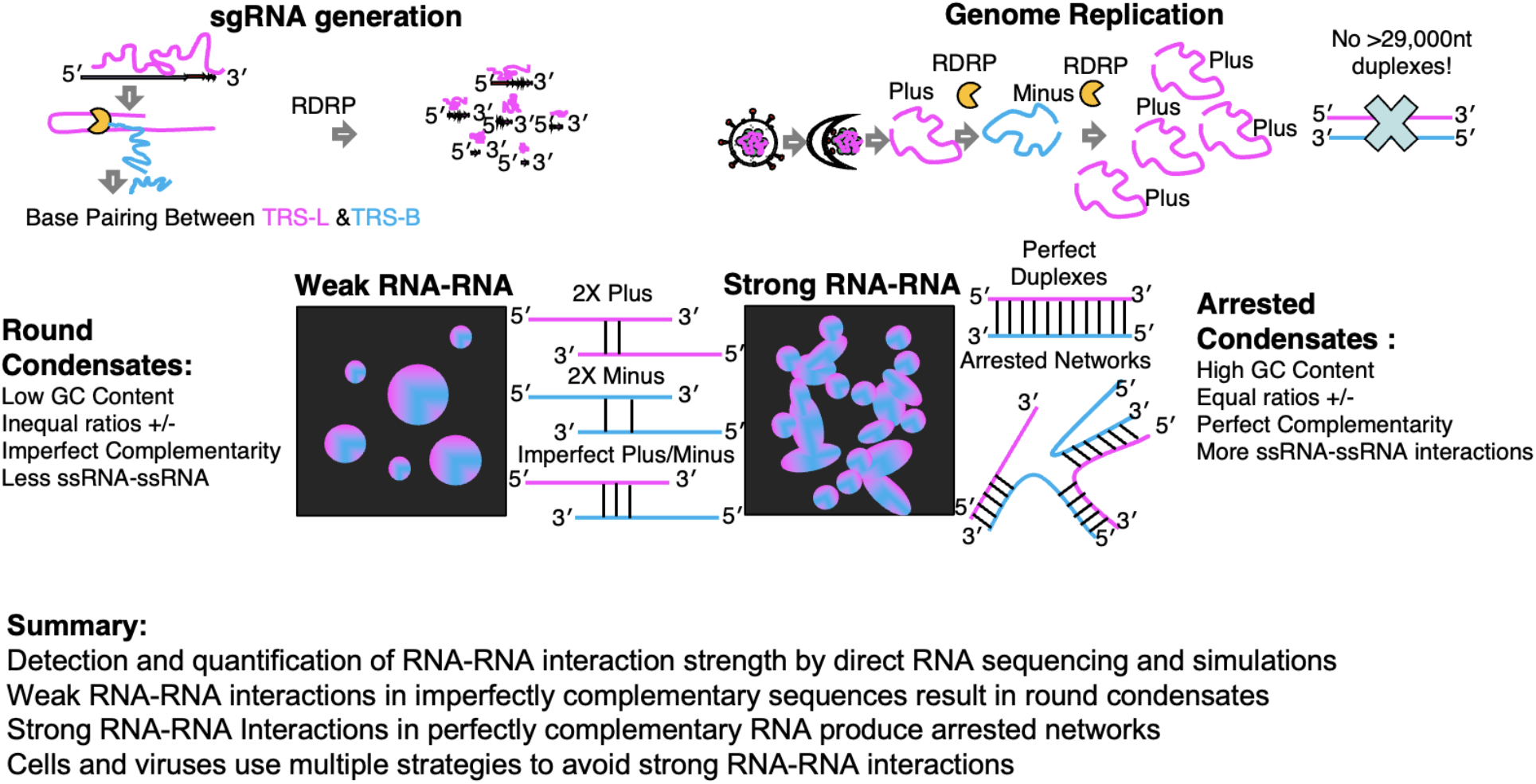

## INTRODUCTION

Proponents of the RNA world hypothesis for the evolution of life suggest that in the primordial soup of ancient earth, the first self-replicating biological polymer was an RNA molecule^1^. In this model, the ability to synthesize a single RNA-binding protein from this self-replicating RNA template was eventually acquired, followed much later by the switch from RNA to DNA as the primary mode of information storage for life. What events would lead to the switch from the purely self-replicating RNA system to self-replicating RNA that synthesizes its own RNA-binding protein?^2^ Two possible models for the selective advantage that could have been conferred by the creation of RNA-binding proteins have been proposed: **(1)** the improved protection of the RNA sequence from the harsh conditions of ancient earth and **(2)** the improved catalytic rate of the self-replicating RNA templates through increased local concentration in the presence of RNA-binding proteins (RBP). These primordial RBPs may have promoted formation of biomolecular condensates^3^, consisting of non-membrane bound assemblies of the self-replicating RNA sequence and its encoded RNA-binding protein. An additional problem in this primordial molecular context is the risk of sense RNA sticking to anti-sense template strand RNA which would inhibit both RNA self-replication and further protein synthesis. We explore here a third possible role for the function of original RBPs in preventing non-productive RNA-RNA interactions.

Any self-replicating RNA polymers of ancient earth are now long degraded, but plus strand RNA viruses provide an extent situation where sense and antisense RNAs must coexist by necessity and at potential peril for RNA entanglement.

Plus strand RNA viruses do not have any DNA phase in their replication cycle and instead generate new copies of their protein-coding capable genomes by creation of a perfectly complementary RNA template sequence consisting of minus strand or anti-sense RNA^4^. Thus, plus strand RNA viruses are thought to resemble the original self-replicating RNAs of ancient earth. As a consequence of this replication strategy, plus strand RNA viruses must produce RNA that is both perfectly complementary and the entire length of the genome. How then do viruses prevent or resolve disastrous genome-long complexes of plus and minus strand RNA, which would prevent protein production and trigger innate immune surveillance via the formation long duplexes that resemble DNA duplexes in length?

To examine how RBPs may chaperone RNA-RNA interactions and mitigate risk of irresolvable and risky RNA duplexes, we study complementary RNA sequences that would coexist and at least transiently interact for SARS-CoV-2 replication. We show that highly stable RNA-RNA interactions produce arrested networks of the RBP Nucleocapsid protein forming condensates. This is mitigated by staggering the addition of the condensing protein (plus, protein, minus), the order observed in the replication strategy of the virus. Thus, RNA viruses may employ an ancient method of coating their genomes in condensing RNA-binding proteins to prevent plus and minus strand RNA–RNA interactions during replication. Our data will show that the formation of biomolecular condensates represents an evolutionary trade off with low affinity interactions supporting beneficial functions and spherical condensate morphologies associated with more dynamic exchange. Conversely, detrimental effects are conferred by high affinity RNA–RNA interactions yielding arrested, gel-like, morphologies. Our data will further demonstrate viruses have likely evolved multiple strategies to mitigate detrimental RNA–RNA interactions and that host eukaryotic cells avoid the problem with DNA genome organization that is depleted in perfectly complementary RNA sequences. Thus, we propose that an original purpose of RNA-binding proteins may have been to prevent spurious RNA–RNA interactions between perfectly complementary strands of RNA and that the innovation of DNA was an attempt to avoid the problem entirely. These data are thus of outsized importance to RNA viruses but likely these principles extend to all RNAs, and RNA encoded biomolecular condensates in cells.

## RESULTS

### Choice of the model biomolecular condensate: Plus strand viral RNA model system

In this study, we are examining the cause and consequence of RNA–RNA interactions in biomolecular condensates. We are testing the two hypotheses that 1) beneficial low affinity interactions are chaperoned by condensates and 2) detrimental high affinity interactions are avoided by the employment of condensing RNA-binding proteins. Both of these can be tested by using the innate replication needs of viruses where plus genome and minus strand templates must be prevented from sticking to each other. By far the most extreme example of this potential problem is found in nidoviruses, which have the longest described genomes of any RNA virus^5^. Nidovirus is the order that encompasses SARS-CoV-2, the virus responsible for the Covid-19 pandemic^6^. Like most plus strand RNA viruses, SARS-CoV-2 encodes its own >29,000 nucleotide long plus strand RNA genome, an RNA-dependent RNA polymerase complex^7^, and an RNA-binding protein capable of undergoing biomolecular condensation with the viral RNA genome, the nucleocapsid protein^8–19^. As a result of extensive pandemic related research, SARS-CoV-2 biomolecular condensates are among the best characterized with respect to the contributions of RNA^11,20–22^, where distinct RNA sequence and structure features yield emergent material properties in the resulting biomolecular condensates. These studies lead to the prediction that specific RNA sequences are in part selected for their impact on specific condensate material properties.

For this manuscript, we will focus on one critical proposed function for SARS-CoV-2 biomolecular condensates, the generation of sub-genomic RNA via RNA–RNA interactions which we and others have proposed could be chaperoned by biomolecular condensate^23–25s^. Utilizing this system, we will first explore whether 5′UTR containing RNA sequences (5′end), plus and minus strand RNA TRS-B sequences, can co-condense.

The name nidoviruses comes from the Latin word for nested, referring to the organizational strategy for the viral RNA genome protein-coding sequences^6,26^. Nidoviruses employ two strategies for protein production. Non-structural proteins (e.g., do not contribute to virion assembly) are generated via a combination of ribosomal frameshifting and protease digestion^27–32^. Structural proteins are generated via the production of sub-genomic RNA (sgRNA)^28,33^. SgRNA production begins during minus-strand RNA production. As the RNA-dependent RNA polymerase synthesizes minus RNA from the 3′ orientation, the synthesis skips from the transcription regulatory sequence body (TRS-B) to the TRS leader (TRS-L) located in the 5′UTR of the genome. The resulting completed minus strand RNA template is used to produce plus strand RNA with one sequence each coding for Spike, ORF3a, Membrane etc., proteins^34^. Productive sgRNA generation is thought to require base pairing between TRS-L (plus) sequences and nascent anti-sense TRS-B sequences (minus)^34–36^. We previously observed that TRS-L/B and TRS-like sequences (YRRRY; where Y = C/U and R = A/G nucleotides) are important drivers of SARS-CoV-2 biomolecular condensation with N protein and are recognized by the N-terminal RNA-binding domain of this protein^25^. In our model, we speculated that the local increased density of N protein condensation promoting motifs on plus strand RNA may help the plus strand RNA genome fold back on itself prior to the skip in minus strand sgRNA production, effectively pinning the complementary sequences of 5′UTR TRS-L and plus TRS-B, nascent minus TRS-B together in the confined volume of the condensate. We further speculated whether such pinning behavior could promote skipping and lead to observed imbalance in sgRNA production correlated with condensation promotion.

### Multiple plus and minus strand TRS-B RNA fragments are capable of driving N protein condensation

To test if biomolecular condensates could promote critical RNA–RNA interactions, we utilized our minimal reconstitution system model of SARS-CoV-2 biomolecular condensates consisting of the structural nucleocapsid protein (N protein) and viral RNA fragments^21^. To this end, we sought to reconstitute the essential TRS-L/B RNA–RNA interaction of nidoviruses.

Thus, we synthesized 13 RNA fragments from the virus that are each 483 nucleotides long, including the 5′end of the genome (TRS-L containing), as well as six regions flanking the TRS-B’s of the Spike, Envelope, Membrane, ORF7, ORF8, and Nucleocapsid RNA in both the plus and minus strand context roughly 237 nucleotides before and after the TRS-B sequence **(Figure 1A**).

**Figure 1.**
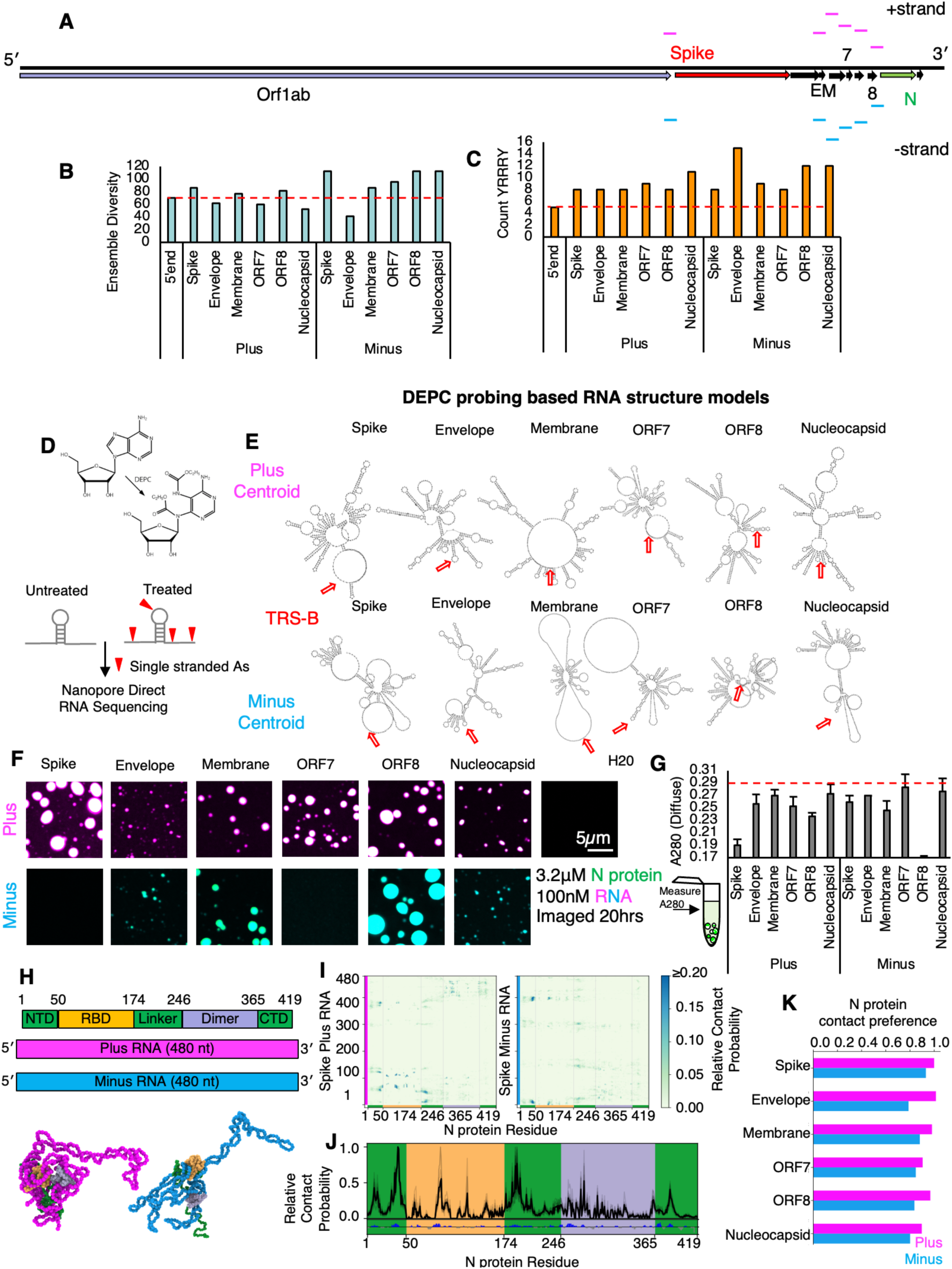
Plus and minus strand TRS-B fragments can undergo condensation with N **protein.** **(A)** Approximate location of tested fragments in the SARS-CoV-2 genome. Plus strand (magenta) minus (teal), spike (red), N/nucleocapsid (light green). Of note, minus strand RNA position is reflective of the complement rather than the reverse complement for ease of comparison to plus strand RNA positions. **(B)** Ensemble diversity predictions for each tested fragment relative to a previously identified condensate promoting sequence, the 5′end. **(C)** Count of YRRRY motifs (N protein N-terminal RNA-Binding domain target sequence) in tested fragments relative to previously identified condensate promoting sequence the 5′end. Of note, this is the total count and is not reflective of local sequence enrichment around TRS-Bs as depicted in **Supplemental Figure S1A**. **(D)** Cartoon depicting RNA structure probing via direct RNA sequencing protocol. Single stranded A nucleotides are preferentially reactive with DEPC (Diethyl pyrocarbonate) resulting in adducts detectable as mutations in nanopore direct RNA sequencing. **(E)** DEPC based RNA structure models for TRS-B (red arrow) containing fragments in plus (top row) and minus Strand (bottom) context. Plus strand RNAs tend to be more structured, particularly around the TRS-B sequence (red arrow). **(F)** Representative images of condensates formed from 100nM RNA TRS-B sequences and 3.2μM SARS-CoV-2 nucleocapsid protein (N protein). Condensates were imaged 20 hours post mixing at the glass. Images reflect a merged signal of RNA (magenta plus and cyan minus) and protein (green) labeled signals. All tested fragments other than spike minus and ORF7 minus resulted in reproducible observable condensates under these conditions. H2O water only no RNA added control indicative of the requirement of RNA to drive N protein condensation under the tested conditions. **(G)** Diffuse phase absorbance measurements for panel D. A280 signal is reflective of the non-condensate recruited portion of the RNA and protein signal. Plus Spike and minus ORF8 are best able to recruit protein and RNA into the dense phase as evidenced by the reduction of signal into the diffuse phase. Error bars are from 3 technical replicates. **(H)** (Upper Panel) Sequence composition of the N Protein and RNA used in simulations. There are 5 domains for N protein: N terminal domain, RNA-binding domain 1, Linker, RNA-binding domain 2, and C terminal domain. For representation of the protein and RNA, we use a hybrid Mpipi-SIS-RNA model (model 1; see Methods). (Lower Panel) Snapshots of dimers of N protein and Spike RNA plus (magenta) and minus (cyan) strands. **(I)** Contact maps of N protein with Spike plus RNA and minus RNA from dimer simulations. **(J)** (Upper panel) Normalized RNA contact probability distribution along the Nucleocapsid protein sequences from all plus and minus RNA sequences in this study. Each histogram from individual simulations is shown as light gray lines, and the overall average is shown as black lines. (Lower panel) The net charge per residue (NCPR) distribution of N protein. The positive charge distribution is highlighted by blue histograms while negative charge is shown in gray. **(K)** Preferences of binding for N protein and RNA from dimer simulations. Higher frequency RNA contacts in plus strand RNA than minus in simulations.

We confirmed that the synthesized RNA fragments that were chosen had a reasonable enrichment of our previously established N protein condensing features^11^. Specifically, we see low ensemble diversity^37,38^ which is indicative of structured/stably folded RNA sequences and promotes the condensation through RNA-binding domain 2 of N protein (**Figure 1B**) and the presence of YRRRY sequences, which promote the condensation through RNA-binding domain 1^25^ (**Figure 1C**, **Supplemental Figure S1A**). These RNA structure and sequence features are comparable to the 5′end RNA sequences that have been previously tested (**Supplemental Figure S1B, C**). To confirm RNA structure predictions, we performed RNA structure probing via direct sequencing^39^ (**Figure 1D and E**). The majority (10/12) newly tested TRS-B fragments were capable of driving condensation with N protein on their own (**Figure 1F**) with 2 minus strand sequences not eliciting N protein condensates. All tested plus fragments drove condensation and, in most cases, also larger assemblies than the minus fragments, consistent with the higher local density of YRRRY in the plus strand (**Supplemental Figure S1A**) and more stable RNA structures (**Figure 1D and E**).

We compared the dilute phase A280 absorbance signal to get “bulk” reaction measurements, following 20 hours of incubation with N protein (**Figure 1G**). Consistent with the observation bulk depletion measurements in **Figure 1G**, minus ORF8 and plus Spike had the largest observed condensate area (**Supplemental Figure S1D**) as well as more protein (**Supplemental Figure S1E**) and RNA signal (**Supplemental Figure S1F**) in the dense phase, these wells had the lowest A280 signal in the diffuse phase (**Figure 1G**) suggesting for these wells, most of the RNA and protein is condensate recruited. To compare the dense phase material properties and the environment, for each RNA system, we quantified the maximum intensity of the protein (**Supplemental Figure S1E**), and the RNA signals (**Supplemental Figure S1F**) in the dense phase of the resulting condensates. Maximum intensity was chosen as it avoids non-uniform loss of signal at the condensate interface (smaller condensates have a larger proportion of interface) and thus is more representative of the densest dense phase environment. To examine the chemical environment, we compared the protein to RNA ratio (**Supplemental Figure S1G**). We observed that, although different total RNA and protein signal was observed in the dense phase depending on RNA sequence, the ratio across tested RNA fragments was quite similar suggesting that each tested RNA fragment should have similar dense phase chemical environment despite differences in condensate size.

To gain additional biophysical insight into the formation of N protein–plus or –minus strand condensates, we conducted explicit chain molecular dynamics simulations. Here, we have implemented a chemically specific residue-resolution approach to probe sequence-dependent binding between the plus or minus strands with N protein and investigate whether these interactions may be sufficient to sustain condensates. Specifically, we use the Mpipi model^40^, a residue-level approach for probing the phase behavior of proteins and RNA. In the original Mpipi implementation, RNA lacks the ability to form base pairs between nucleotides. Thus, we updated the RNA representation with the RNA-SIS model, a consistent nucleotide- resolution model^41,42^ that enables explicit RNA base pairing. In the resulting protein–RNA model (referred herein as model 1; see Methods), each amino acid or nucleic acid is represented as a single interaction site, disordered protein regions as fully flexible chains, folded regions of N protein via homology-modeled structures, and nucleotides can explicitly form base-pair interactions (hence, can stabilize secondary structures). Additionally, proteins and RNA interact via both non-charged and charged interactions, where interactions of amino acids with G/A nucleotides are favored over those with C/U based on atomistic predictions. All simulations with model 1 were performed in the NVT ensemble at 300 K and 150 mM NaCl salt. Further model and simulation details are included in the Methods section.

We first simulated dimeric systems, composed of N protein and plus or minus strands. As shown in **Figure 1H**, there are five regions in N protein: the N terminal domain (NTD), RNA binding domain (RBD) 1, Linker, RBD 2, and C terminal domain (CTD). Among these regions, NTD, linker and CTD are disordered (green), while the RNA binding domains are mainly folded.^15^ In our dimer simulations, all plus and minus strands bind N protein. However, different regions on plus and minus strands show high residue–residue contact frequencies with N protein (**Figure 1I, Supplemental Figure S1H**). As an example, the contact map of N protein with Spike plus and minus RNAs are depicted in **Figure 1I**. To further investigate the binding mechanism, we assess normalized contact probabilities between all RNA strands and N protein (**Figure 1J**). Despite the sequence variations, the nonspecific binding pattern, with respect to N protein, is highly preserved among the RNA strands. The charge pattern analysis of N protein reveals that the observed nonspecific binding patterns are mainly governed by the charge distribution and the structure of N protein (**Figure 1J**). The positive charge distribution highly correlates with the peaks of contact probability. Interestingly, even though the positive charges are almost evenly distributed throughout the sequence of N protein (**Figure 1J**), the high-probability binding sites are mainly concentrated around RBD 1 and its neighboring disordered regions—suggesting an importance of protein structure for this binding interaction. Moreover, our findings for full-length N protein with long RNAs align well with previous studies, where the dimer of partial N protein (NTD+RBD 1) and short RNAs (poly rU) were studied using Mpipi model.^43,44^ Here, we predict that for the full-length N protein with much longer RNA strands, the NTD–RBD emerges as the dominant motif for interaction with RNA. Furthermore, due to size difference between the N protein and RNA, only part of the RNAs bind to N protein. As a result, the unbound nucleotides are free to base-pair with other RNA molecules or interact with additional N proteins. These data suggest that both plus and minus strands bind N protein and that non-specific charge-dominated interactions underlie the formation of condensates observed in the experiments (**Figure 1F**). We examined the preferences of binding for N protein and RNA from dimer simulations by quantifying the contact frequency in tested RNAs (**Figure 1K**). In keeping with our results from **Figure 1F**, plus strand RNA regardless of sequence had higher affinity for simulated N protein. As noted earlier, plus strands are enriched in YRRRY sequences that facilitate condensate formation via interactions with N protein RBD 1. Given that our model encodes stronger protein–RNA interactions for the larger G/A nucleotides, the simulations effectively capture such non-specific interactions between YRRRY sequences and the RBD 1.

However, in our simulations we do not observe a dominance interaction between RBD 2 and the more structured plus strands. We believe that this may likely be a limitation of the model in capturing more specific interactions between proteins and RNA.

With this information we next sought to test combinations of plus and minus RNA TRS-B fragments with the 5′end (**Figure 2**) to reconstitute a critical viral RNA–RNA interaction.

**Figure 2.**
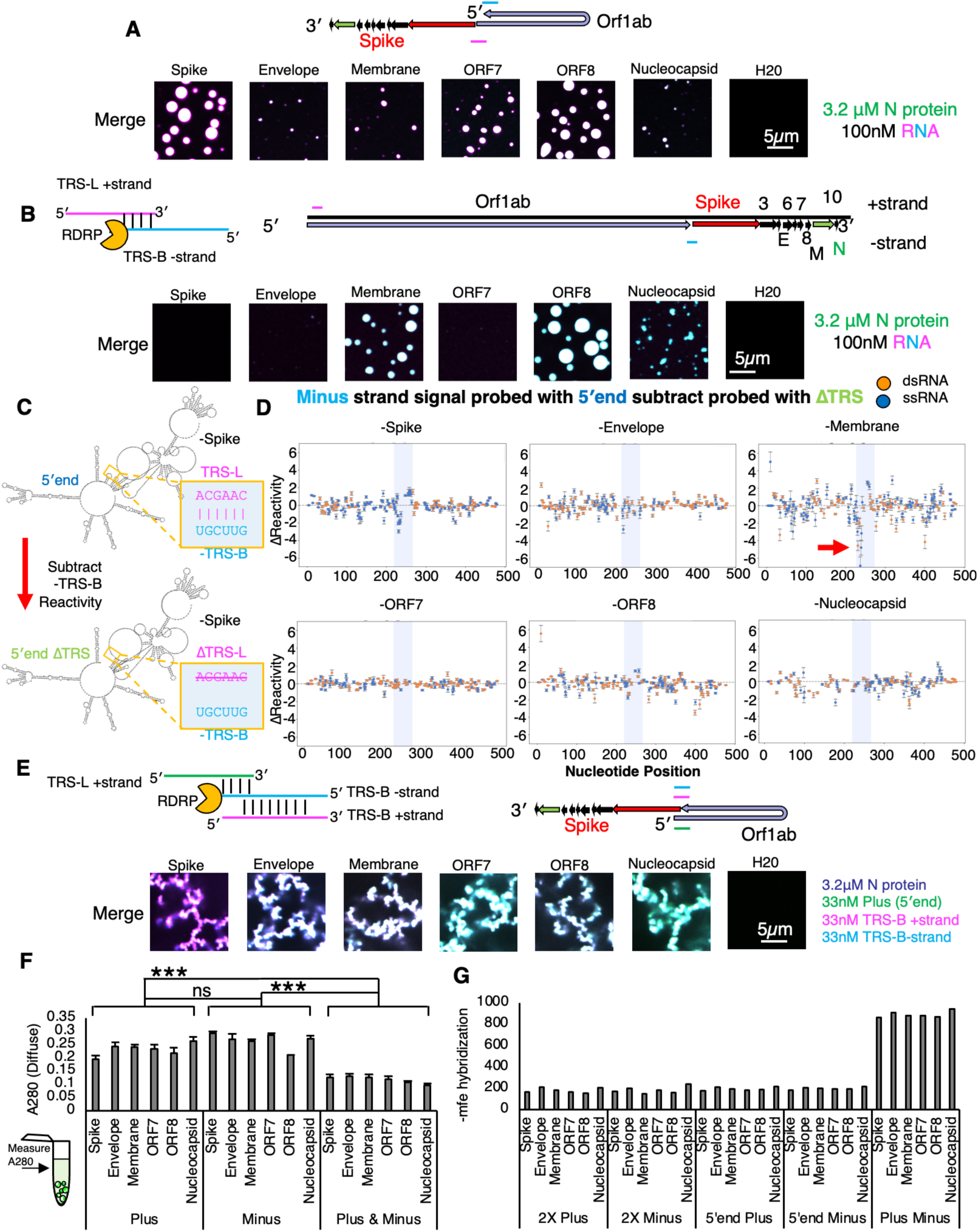
Reconstitution of 5′end RNA looping and sgRNA generation 5′end RNA co-condenses with TRS-B fragments. For **A, B**, and **E**, all images depict the merged signal of the same total RNA (100nM) and protein (3.2μM) concentrations. As shown in panels **2A** and **2B** combinations of two RNAs are 50nM each for panel **2E** combinations of RNA are 33.3nM each. All images were acquired 20 hours post mixing at the glass. Scale bar (white line in H_2_O only control) is 5 microns. All images in each panel are acquired on the same day and are contrasted the same. **(A)** Cartoon depicts the theoretical arrangement of the plus strand RNA genome to tether the 5′UTR in proximity of the TRS-B of spike (red) prior to minus strand RNA synthesis which may be assisted by condensate formation. Combinations of 5′UTR containing RNA (5′end labeled in cyan) and plus TRS-Bs labeled in magenta. Depicted images are representative of merged signal (N protein in green and RNA in cyan and magenta) of 3 technical replicates and illustrate that 6/6 tested TRS-B containing plus RNAs can co-condense with 5′end in the N protein condensates. Unmerged images are depicted in **Supplemental Figure S2A**. H2O is water only no RNA control. **(B)** Cartoon depicts the theoretical arrangements base pairing between the plus TRS-L containing 5′end and minus strand TRS-Bs prior to sgRNA generation which may be assisted by condensate formation. Base pairing as indicated by black lines is indicated between the plus TRS-L and minus TRS-B. Genome position of minus fragments is reflective of complement rather than reverse complement positions. Combinations of 5′UTR/TRS-L containing RNA (5′end labeled in magenta) and minus TRS-Bs labeled in cyan. For minus TRS-B RNAs capable of driving condensation, 5′end and minus can co-condense. Depicted images are representative of merged signal (N protein in green and RNA in cyan and magenta) of 3 technical replicates and illustrate that RNAs can co-condense in the condensates. Unmerged images are depicted in **Supplemental Figure S2B**. H2O is water only no RNA control. **(C)** Overview of experimental protocol to detect RNA–RNA interactions in trans. Two RNA species are mixed in equal molar ratios (e.g. 5′end and minus strand RNA e.g. - spike) if base pairing occurs (pink and blue sequences) a change in the single stranded reactivity in the minus should be evident particularly when the sequence is absent in the plus (e.g. co-incubation with TRS-Del RNA). **(D)** Δ Reactivity plots for each minus RNA representing the change in signal due to the presence or absence of co-incubated TRS-L containing sequences in the 5′end. Minus membrane RNA shows a large decrease in A reactivity near the TRS-B (blue box/red arrow) when TRS-L is present as opposed to absent ΔTRS-L indicative of base pairing between the TRS-L and B. Change in RNA signal is most easily observed in single stranded A nucleotides (blue circles ssRNA) as opposed to double stranded A nucleotides (orange circles dsRNA). **(E)** Cartoon depicts the theoretical arrangement of plus RNA genome (plus TRS-L and plus TRS-B) as well as minus nascent synthesized RNA genome just prior to sgRNA synthesis. Base pairing as indicated by black lines could take place between TRS-L and minus but should be more extensive between minus and plus TRS-B containing fragments. Combinations of 5′UTR containing RNA (5′end labeled in green) and plus TRS-Bs in magenta minus TRS-Bs labeled in cyan. Of note a difference from all previous protein labels N protein is labeled in blue (Atto405) rather than green. Depicted images are representative of merged signal of 3 technical replicates and illustrate that RNAs can co-condense in the condensates. In contrast to A and B, condensates no longer adopt a rounded morphology. Unmerged images are depicted in **Supplemental Figure S2C**. H2O is water only control (no RNA). **(F)** Diffuse phase absorbance measurements for panels A, B and E. A280 signal is reflective of the non-condensate recruited portion of the RNA and protein signal. Despite containing identical concentrations of RNA and protein, combination which contain 3 RNAs result in more recruitment to the dense phase and lower diffuse phase signal whereas 2 RNAs were not significantly different from each other by this metric (p<0.001 ***) **(G)** Prediction of RNA hybridization energy between any 2 RNA species tested thus far (2× plus and 2× minus) (Figure 1F), 5′end/plus (**2A**/**E**), 5′end minus (**2B**/**E**), and plus/minus (**2E**). Most RNA combinations have a low -MFE prediction indicative of poor RNA–RNA interaction. Plus/minus has a high predicted -MFE indicative of strong RNA–RNA interaction. Of note, -MFE is predicted rather than MFE as positive numbers can be more intuitive.

### Reconstitution of low and high affinity viral RNA–RNA interactions in condensates

During sgRNA generation, the RdRp jumps from the TRS-B to the TRS-L sequences skipping ORF1ab sequences in a base pairing and N protein dependent process^27,31,34^. We have postulated that N protein/TRS-RNA condensation may also help promote this skipping by tethering the two RNA sequence in a smaller volume of the condensate^25^. We reasoned that the comparatively higher local density of N protein condensation promoting motifs on the plus RNA as observed in **Figure 1** and **Supplemental Figure S1A** might allow for co-condensation of the 5′end and the TRS-Bs effectively tethering the 5′end in place adjacent to the location of the jump in cis. This tether, which could arise by N protein condensate formation, may support sgRNA generation by helping the polymerase skip from the TRS-B to the TRS-L, looping out the intervening plus strand genome sequence. To test if the 5′end and the plus TRS-Bs containing fragments could co-condense, we reconstituted labeled 5′end RNA with labeled TRS-B containing fragments. We were able to observe that, for fragments that resulted in condensates, both labeled RNAs were present in the dense phase (**Figure 2A, Supplemental Figure S2A**).

Extensive base pairing between the plus strand 5′end TRS-L sequence and the minus strand TRS-B is thought to be essential for the proper generation of the minus strand sgRNA sequence^33,34^. This is because base pairing may promote skipping of the RNA dependent RNA polymerase from the body to the leader resulting in the needed chimeric sequence. To test if minus strand RNA could also enter co-condensates with the 5′end and thus promote this interaction we tested the labeled 5′end RNA with each of the labeled TRS-B fragments in the minus strand context. Again, we observed under the tested conditions that the primary arbiter of condensation appeared to be the TRS-B minus strand RNA identity and minus strand RNA was poorly able to drive condensation (**Figure 2B, Supplemental Figure S2B**).

To confirm that TRS-L and B sequences were capable of pairing in our experimental conditions and without N protein, we performed RNA-RNA interaction mapping for each of the tested minus TRS-B fragments co-incubated with the plus 5′end fragment which contains either the wildtype or TRS-L deleted sequence. We could identify regions in each minus strand RNA that were paired in trans to the TRS-L by comparing the reactivity of nucleotides when probed in the minus TRS-B RNAs in either the presence of 5′end or the 5′end ΔTRS sequence (**Figure 2C**), By subtracting the probed with ΔTRS-L from the wildtype we observed the greatest change in reactivity was found in the region surrounding the minus TRS-B (light blue box). (**Figure 2D**), suggesting that most of our RNA combinations had successfully reconstituted base pairing between the plus TRS-L and minus TRS-B. Of note, the highest change in reactivity (indicative of stronger base pairing) was observed in the -membrane RNA fragment. Intriguingly, this RNA was the most single stranded (blue dots) in the region surrounding the TRS-B in our RNA structure data (**Figure 1E**) suggesting that the single stranded RNA content in the area surrounding the minus TRS-B may facilitate pairing between the TRS-L and TRS-B.

With this information, we reasoned that if our model was correct and N protein co-condensation was able to promote the jump between TRS-B and TRS-L, all three RNA sequences would be present together in the same location^34^, perhaps in the dense phase of an N protein biomolecular condensate^25^, just after minus strand synthesis. Thus, we differentially labeled all three RNAs and mixed them with labeled N protein. Surprisingly, under the tested conditions, not only were all three RNAs present in the same dense phase, but the morphology of the resulting assemblies also differed significantly from any previously tested condition with condensates consistently forming giant arrested networks (**Figure 2E, Supplemental Figure S2C**) Comparison between the diffuse phase of all three tested conditions suggested that presence three RNAs, despite having the same total RNA concentration (100nM in all cases), was much better at recruiting RNA and protein to the dense phase than single RNAs (**Figure 1F**). We reasoned that the profound morphology change observed in the condensates may be due to a difference in RNA–RNA interaction strength^37,38^. To this end, we predicted the hybridization energy^45^ (**Figure 2G**) of all RNA combinations tested so far double the concentration or 2× plus and 2× minus (**Figure 1F**), plus with 5′end (**Figure 2A**), minus with 5′end (**Figure 2B**), and plus with minus (**Figure 2E**). We observed that the highest predicted hybridization energy was found in the plus/minus RNA combinations consistent with the perfect complementarity of these RNAs. Very similar lower values were observed for every other combination. Thus, we reasoned that the profound change in morphology was well correlated with the strength of RNA–RNA interaction in the condensate (**Figure 2G**).

### Reconstitution of perfectly complementary RNA–RNA interactions alone is sufficient to yield arrested networks

To test if plus and minus strand RNA–RNA interaction strength alone was sufficient to drive the observed morphology change, we next made condensates that contained only these perfect plus/minus RNA pairs (**Figure 3A**).

**Figure 3.**
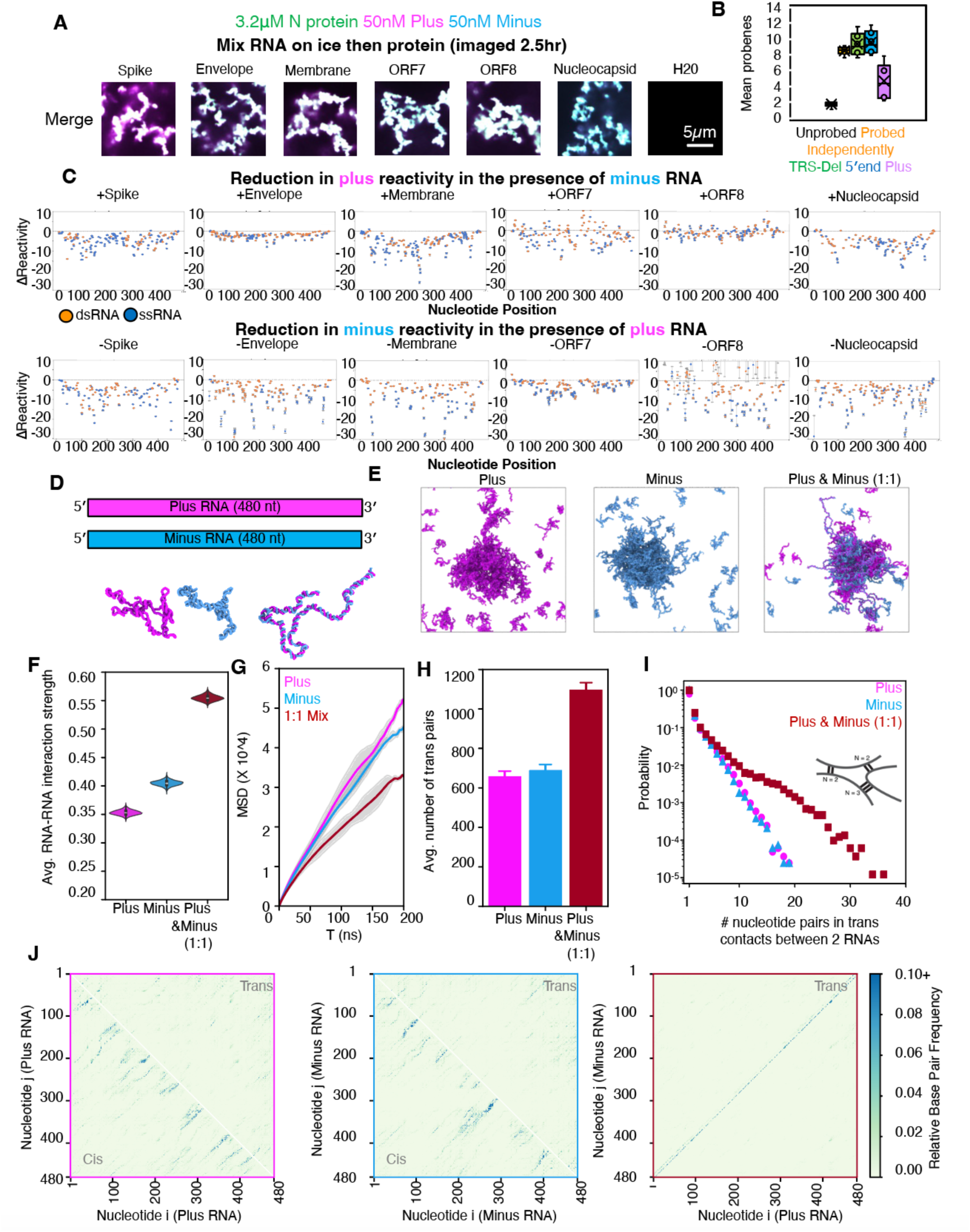
Mixtures of perfectly complementary plus and minus strand RNA result in arrested network formation due to increased RNA-RNA interactions. **(A**) Premixing 50nM each of plus and minus RNA on ice followed by N protein addition (3.2μM) is sufficient to produce arrested networks. For **A**, all images depict the merged signal of the same total RNA (100nM) and N protein (3.2uM) concentrations. Combinations of two RNAs are 50nM each. Scale bar (white line in H_2_O only control) is 5 microns. All images in each panel are acquired on the same day and are contrasted the same. **(B)** Mean probeness of minus strand RNA, without probing (black), probed independent of any plus RNA (orange), probed in the presence of imperfectly complementary RNA (green 5′end ΔTRS, blue 5′end wildtype) or probed in the presence of perfectly complementary RNA (purple). Reduction of probed signal, indicative that base pairing is strongest when minus strand RNA is probed in the presence of perfectly complementary plus RNA. These data are indicative of base pairing in trans in this condition. **(C)** Nucleotide resolution Δ reactivity plots (probe plus/minus subtract probed alone) for plus strand RNAs (top panel) and minus strand RNAs (bottom panel). Non-uniform decrease in signal in the presence of complementary RNA as opposed to independently probed. Orange dots are double stranded A nucleotides and blue dots are single stranded A nucleotides. Greater reduction in reactivity is observed for minus strand RNA and single stranded A nucleotides. **(D)** Representative single chains of perfectly complementary plus (magenta) and minus (cyan) RNA base pair in trans in RNA only simulations (model 2; see Methods). **(E)** Representative images depicting Envelope RNA simulations of 2X plus (magenta), 2X minus (cyan), or 1:1 plus and minus RNA. The analysis for these simulations follows: **(F)** Average RNA–RNA interaction strength. This is obtained by normalizing the potential energy of each system by the total number of nucleotides. **(G)** Comparison of mean-squared displacement (MSD) of center of mass. Here, the 1:1 mixture exhibits the slowest dynamics. **(H)** Average number of nucleotides pairs in trans RNA contacts. **(I)** Probability distribution of number of nucleotide pairs in trans contacts. **(J)** Cis and trans contacts for plus-only (left panel) and minus-only (middle panel) systems, as well as trans contacts for 1:1 mixture (right panel).

We observed that for all tested RNA pairs (6/6) mixing perfectly complementary plus and minus strand RNA were all capable of driving the networked condensates phenotype (**Figure 3A**). To determine if the morphology of the condensates was due to RNA–RNA interaction strength, we measured RNA–RNA interaction strength directly using DEPC probing and sequencing^39^ (**Figure 3B**). We observed unprobed minus RNA, regardless of sequence, had low level of probing as evidenced by a lack of mutant Adenosines in the raw sequencing data.

Minus RNA that was probed independently of any plus RNA had much higher level of probing. Minus strand RNA that was probed in the presence of plus strand RNA that was not perfectly complementary (5′end or TRS-del) had on average comparable levels of reactivity to minus strand RNA that was probed independently of any other sequence being present. In striking contrast, minus strand RNA that was probed in the presence of perfectly complementary plus strand RNA showed a profound reduction in probing signal, perhaps due to trans base pairing of the previously single stranded Adenosines in the minus strand to the complementary Uracils in the plus strand rendering them inaccessible to chemical probing by DEPC (**Figure 3B**). To examine the nucleotide level changes in reactivity, we plotted the Δ Reactivity (the subtraction of the signal for the RNA probed independently from the signal for RNA probed in the presence of its complement) (**Figure 3C**). We observed that individual nucleotide reactivity had differential sensitivity and that plus stranded RNA signal change was far less uniformly reduced than minus. This variability in the sensitivity may be due to differences in single strandedness of the nucleotides in plus versus minus strand RNA. As plus RNA tends to be more structured than minus RNA^46^ (**Figure 1B and 1E**), changes in individual nucleotides are more variable (**Figure 3C**). Of note, and in keeping with previous observations from Trcek lab^47^, single stranded RNA/ssRNA reactivity signal (blue dots) was more altered than double stranded RNA/dsRNA indicative of more base pairing for single stranded RNA in the presence of its RNA complement. RNA–RNA interaction mapping by sequencing gives us nucleotide and single molecule resolution of trans RNA–RNA contacts. It does not however give us a 3-dimensional arrangement of the RNA molecules in contact in the dense phase. To examine the arrangement of RNA molecules, we turned to molecular dynamics simulations of the RNA alone. This approach aimed to better understand the molecular origins for the arrested phenotype observed for N protein–plus–minus systems, as all our data up to this point indicated that the condensate morphology change was primarily RNA–RNA interaction strength dependent.

We hypothesized that an enhanced interaction strength from perfectly complementary RNA–RNA interactions could lead to kinetically arrested states, independent of N protein–RNA binding (and consistent with our RNA only sequencing data from (**Figure 3B and C**). To test this, we adopted a minimal RNA model^41,42^ (model 2; see Methods) that describes RNA–RNA binding based on canonical and wobble base-pairing propensities. The model is trained to capture both cis and trans RNA–RNA contacts in the presence of high salt (i.e., effectively neutralized backbone charges) and, therefore, captures RNA clustering in the absence of protein (**Figure 3D, E**). Employing this approach, we investigate the clustering of plus and minus strands individually, as well as mixtures of the two strands (**Figure 3E, Supplementary Figure S3A-C**). In all cases, our simulations predict stronger RNA–RNA interaction strength for RNA clusters with perfectly complementary RNA–RNA interactions compared to their plus/minus strand analogues (**Figure 3F, Supplemental Figure S3G)**. Additionally, the degree of RNA–RNA interactions was directly but not perfectly proportional to the GC content of the RNA (*R*^2^ = 0.61) (**Supplementary Figure S3K and L**), suggestive of sequence and structure mediated effects on RNA–RNA interaction. Another common trend (except ORF7) is that plus-strand clusters in general have lower RNA–RNA interaction strength than the minus strands.

This also might be inherent to plus strands, which need to be less prone to form aggregates for a functioning virus replication cycle. Furthermore, as mentioned above, the plus strands are more structured than their minus strand counterparts (**Figure 1E**). This suggests that plus strands are overall less free to base-pair, which is consistent with their lower RNA–RNA interaction strength in our simulations (**Figure 3F**). Interestingly, the reactivity for ORF7 shows an opposite trend to the other sequences (**Figure 3C**), which is also mirrored by the simulations (**Supplemental Figure S3G**).

Next, we quantified the diffusivity of RNA strands in each of the systems to investigate if perfectly complementary RNA–RNA interactions lead to RNA clusters with reduced dynamics. Our analysis revealed that, on average, RNA strands are less dynamic in systems with perfectly complementary RNA–RNA interactions (**Figure 3G, Supplemental Figure S3H**). Notably, the diffusion exponent here is less than 1 for all simulated systems, indicating sub-diffusive behavior of RNAs without the presence of proteins. Therefore, N protein in protein–RNA-condensates maintains liquid-like properties by mediating extensive RNA pairing by its binding to RNA (**Figure 1H-J**) or by other mechanisms.

To gain microscopic insight into the trends in RNA–RNA interaction strengths and RNA strand dynamics, we quantified the degree of trans RNA contacts in our simulations (**Figure 3H**). Indeed, the 1:1 mixture with perfectly complementary RNA–RNA interactions have roughly two-fold greater trans RNA contacts than plus- or minus-strand only systems (**Figure 3H, Supplemental Figure S3I**). Additionally, the probability distribution of nucleotide pairs in trans RNA contacts is right-shifted for the systems with perfectly complementary RNA–RNA interactions (**Figure 3I, Supplemental Figure S3J**), indicative of extensive trans RNA contacts. Furthermore, plus- and minus-strand only systems reveal more randomized contact pattern at the nucleotide level (**Figure 3J**). In contrast, the 1:1 mixture exhibits a highly ordered nucleotide contact pattern, consistent with the perfectly complementary nature of the plus and minus strands. Thus, these results strongly suggest that the arrested networked condensates primarily arise from extensive trans RNA contacts rather than random RNA entanglement.

Together, our experiments and simulations suggest that perfectly complementary RNA– RNA interactions could lead to an arrested phenotype during viral replication.

### Coating with N protein reduces trans RNA-RNA interactions

Given the strong impact duplex RNA has on arresting condensates, how do viruses resolve the RNA duplexes that are an unavoidable step in the replication cycle of RNA viruses (in the case of SARS-CoV-2 >29,000 nucleotides long)? We hypothesized that separation in timing between protein binding and replication of the genome may be a mechanism to prevent the potentially deleterious duplexes. In the case of SARS-CoV-2, one complete plus strand copy of the genome enters the cell which could be bound by the ∼1000 N proteins which also enter with the virion^48^. From this single plus strand RNA, a minus strand is synthesized (**Figure 4A**). Thus, binding/condensation with N protein or other RNA binding proteins might prevent the sticking of the plus strand RNA to the minus strand RNA pre-coating plus strand RNA with protein. We reasoned we could mimic this experimentally by altered timing of addition of RNA to protein.

**Figure 4.**
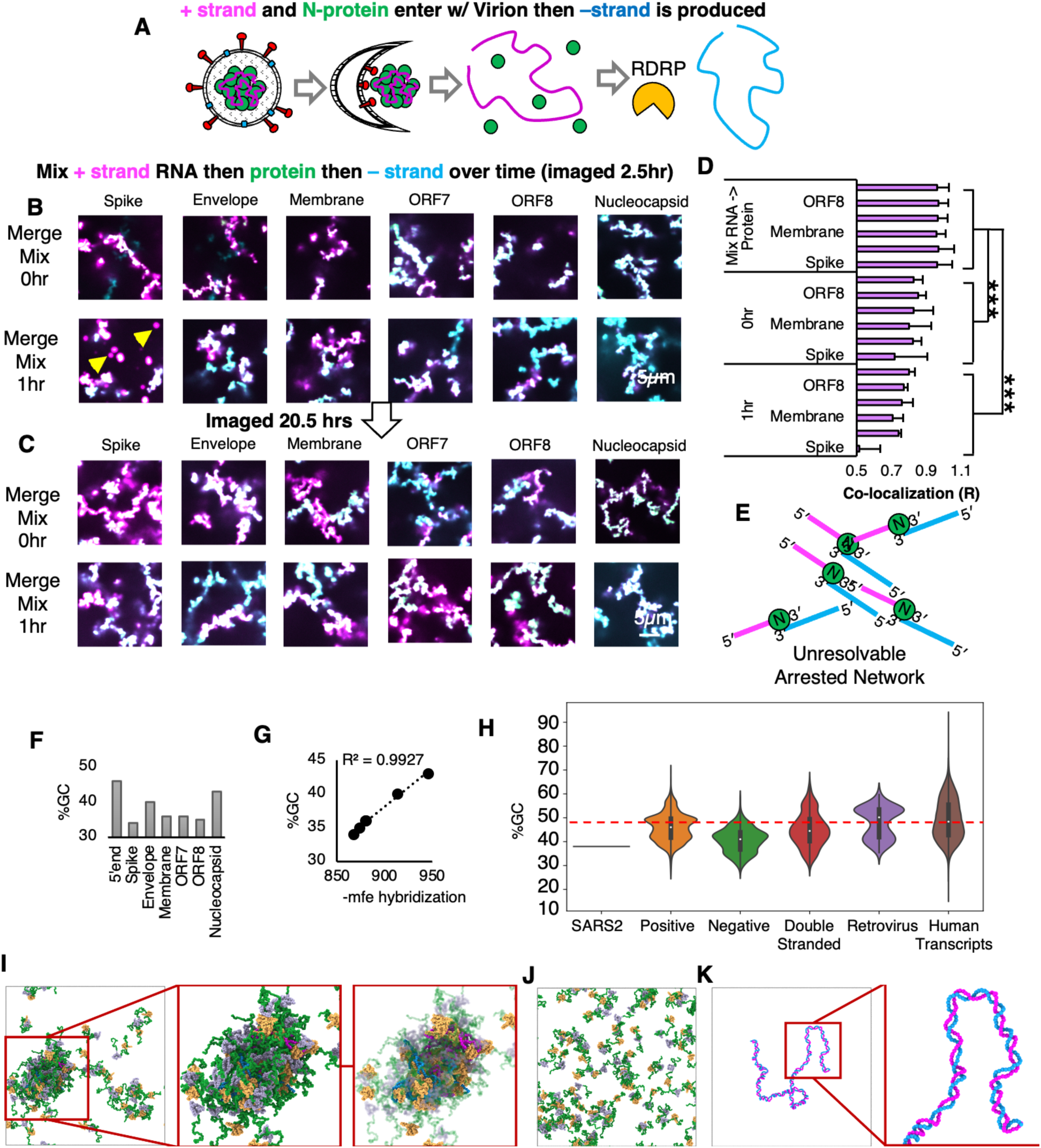
RNA addition timing and GC content modulate RNA-RNA interactions **(A**) Cartoon depicts the genome replication cycle of SARS-CoV-2, applicable to many plus strand RNA viruses. Plus stranded RNA and N protein precedes minus strand RNA production. Plus strand RNA (pink squiggle) enters the cell, bound by N protein (green dots) in ribonucleoprotein (RNP) complexes (30-35 per virion for SARS-CoV-2). Upon entry into the cytoplasm, N protein uncoating (likely by ribosome read through) allows for the translation of multiple proteins including the RNA-dependent RNA polymerase (RDRP orange Pacman). RDRP uses the plus strand RNA genome as a template to produce the minus strand RNA (teal squiggle). For Figure 4B and C, all images depict the merged signal of the same total RNA (100nM) and N protein (3.2μM) concentrations. As shown in panel Figure 3A combinations of two RNAs are 50nM each. Scale bar (white line in H_2_O only control) is 5 microns. All images in a given panel are acquired on the same day and are contrasted the same. **(B)** Staggering the addition of minus RNA to plus RNA and protein by 0, and 1 hour prevents complete mixing of plus (magenta) and minus (cyan) RNA signals. Of note at this timepoint for 1 hour post preincubation, some amount of plus Spike RNA is still in spherical assemblies (magenta circles as marked by yellow arrows). **(C)** Related to **B** additional incubation following preincubation does not lead to increased co-localization in the RNA channel (plus magenta minus cyan mixing for any tested RNA combination. Quantification in **Supplemental Figure S4C**.) **(D)** Quantification of the reduced co-localization of plus and minus RNA signals following preincubation of plus RNA (Figure 4A) with N protein as compared to pre-mixing of RNA on ice (Figure 3A). In all tested RNA combinations, for all 3 technical replicates of each condition (error bars), preincubation results in a reduction of colocalization (R) whereas premixing, R values are close to 1 indicating near perfect co-localization of plus and minus RNA signals. **(E)** Model of the arrangement of plus and minus strand RNA molecules with N protein condensates which could result in an unresolvable arrested network morphology. **(F)** GC content of tested TRS-B fragments with Spike TRS-Bs having the lowest %GC and Nucleocapsid having the highest. (Of note plus and minus strand RNA of the same type e.g. spike has identical GC content). **(G)** %GC content is correlated with -mfe of RNA–RNA interaction strength, indicating that the strength of RNA–RNA Interaction is proportional to the GC content with plus/minus spike being the least “sticky” and plus/minus nucleocapsid being the most “sticky”. Consistent with this, predicted spike RNA has the least propensity to be mixed at 2.5 hours panel **4C** and **4D**. **(H)** Genomes of RNA viruses are depleted in GC content compared to the human transcripts. SARS-CoV-2 has very low fraction of GC content compared to human transcripts and plus strand RNA viruses in general. Negative stranded RNA viruses have the lowest GC content on average. Red dotted line depicts 50% GC content.) **(I)** Coarse-grained simulation of an N protein–RNA system (64 N protein; one plus and one minus Envelope RNA). Simulations are performed using model 1 (see Methods). Zoomed in snapshots of the protein–RNA cluster are shown. For clarity the transparency of portions of proteins in the rightmost snapshot are enhanced to reveal the protein coating on RNA. **(J)** Coarse-grained simulation (model 1) of a pure N protein system (no RNA). Without RNA, N protein does not form clusters. **(K)** Coarse-grained simulation of one plus and one minus Envelope RNA strand, using model 1. In this case, two RNA strands form a perfectly complementary dimer that is sustained over a one microsecond simulation.

To test the hypothesis that timing of minus strand RNA addition relative to protein might mitigate strong RNA–RNA interactions we altered the timing of addition of RNA with plus RNA, N protein, and minus RNA—representative of the proposed pre-binding of N protein to the plus strand prior to the production of minus by the RNA-dependent RNA of polymerase. To this end, we first mixed N protein to plus strand RNA and then staggered the addition of minus strand to this mix by 0, and 1 hours (**Figure 4B**). We assessed the ability of N protein to mitigate RNA-RNA entanglement by measuring the degree of colocalization^49^ of the plus and minus strand RNAs in the same pixel (**Figure 4D**). The presence of protein prior to the plus and minus strands encountering one another was sufficient to prevent the plus and minus RNAs from mixing well for all the tested RNA sequences (**Figure 4B-D**). Remarkably even the almost immediate addition (0 hour) of minus RNA did not mix well with plus RNA that has been premixed with N protein, even after 2.5 hours of incubation of all three components together. We quantified the difference between premixing RNA on ice (**Figure 3A**) and staggering minus strand addition after plus/N protein mixing by measuring the co-localization (R) of the plus and the minus strand RNA signals (**Figure 4D**). Diffuse phase RNA measurement suggested that the dense phase recruited different levels of RNA and protein depending on staggering consistent with longer incubation with the higher condensate promoting sense RNAs better able to recruit RNA to the dense phase (**Supplemental Figure S4A**). We observed that for all tested RNA combinations staggering the minus strand RNA addition resulted in a statistically significant reduction in RNA co-localization indicating the prevention of complete mixing between the opposed RNA strands (**Figure 4D**).

Many condensates display liquid-like material properties^50–58^. If these properties were present in our arrested networks, we would predict that additional incubation would allow for the complete or more complete mixing of RNA, with continued failure to mix indicative of more solid or aggregate-like material properties^59^. To determine if continued incubation would allow for additional mixing, we again imaged the condensates 18 hours later (approximately 20.5 hours from the mixing of plus RNA and N protein). We observed that the plus and minus strand RNA signals continued to be not co-localized even 18 hours later (**Figure 4C**) with no significant difference observed between 2.5-hour observation timepoint and 20.5 hours observation timepoint (**Supplemental Figure S4B and C**).

The lack of complete mixing of plus and minus strands suggests that the coating of RNA by N protein, mediated by favorable N protein–RNA interactions, aids in suppressing trans RNA contacts. Interestingly, recent work has predicted a strong interaction motif between N protein and RNA sequences—with the NTD and RBD 1 of N protein forming a positive groove that binds RNA^60^. Thus, we also hypothesized that such an interaction motif could out-compete RNA–RNA binding, effectively reducing sticking of RNAs to one another. To probe this at submolecular resolutions, we simulated systems composed of 64 N proteins and two complementary RNA strands (Envelope), which represents a system with excess N proteins, which is consistent with the ratio of protein-to-RNA in our in vitro experiments. We then assess the formation of protein– RNA clusters in our simulations. Here a cluster is defined using a distance-based metric (see Methods). There are about 37 N proteins and 2 RNAs in the formed RNA–protein clusters, which is a charge balanced case. Interestingly, in early viral infection, it is estimated that approximately 1,000 N proteins colocalize with a 30-kb RNA strand^48^, which is consistent the charge ratio in our predicted protein–RNA cluster. Surprisingly, despite being in proximity, the two RNA strands form no stable trans RNA contacts due to excessive N protein coating (**Figure 4I**). Multiple RBD domains, along with regions of N protein, can coat the same RNA strand, thereby reducing the probability of extensive base pairing. In **Figure 4I**, the RBD 1 coating is easier to detect by making the remaining domains of the proteins semi-transparent. As a reference, in these simulation conditions N protein is not able to form clusters without RNA (**Figure 4J**), and the two complementary RNA strands are able to sustain extensive base pairing in the absence of N protein (**Figure 4K**). Collectively, these simulations support the model that the coating of plus strand RNA by N protein can prevent mixing of the plus and minus strand RNAs. However, exposed non-coated regions in plus might still be able to base pair with perfectly complementary minus strand RNA resulting in the arrested networks observed experimentally (**Figure 4E**). Thus, additional strategies other than coating of plus RNA by N protein prior to minus strand RNA production must be employed to mitigate spurious and potentially strong complementary RNA interactions between the plus and minus strands.

Our simulations and wet lab experiments revealed that coating RNA in N protein reduced spurious interactions and co-localization. We next asked if there were any RNA sequence specific differences in co-localization in our 6 tested combinations. Condensates with longer plus strand only incubation time recruiting more protein to the dense phase (**Supplemental Figure S4A**), consistent with the higher propensity of plus strand RNA for driving N protein condensation but there was no obvious consistent trends with plus strand RNA alone condensates **(Figure 1F and G)** suggesting plus strand affinity to N protein did not play a significant factor in suppression of RNA-RNA interactions by pre-coating with N-protein. Examination of the RNA signal reveal that RNA signals failed to mix upon staggered addition, and the degree of colocalization was not uniform with the lowest degree of mixing as quantified by plus/minus RNA low colocalization score observed in Spike TRS-B fragments (**Figure 4D**), (**Supplemental Figure S4D**). Further, colocalization of RNA signal was not altered with longer incubations times (2 vs. 20hr) suggesting that once arrested networks form they are irresolvable under tested parameters (**Supplemental Figure 4C**) Additionally, when the normalized colocalization signal was compared for the 0-, and 1-hour conditions, only Spike was significantly less mixed (**Supplemental Figure S4D**) following incubation time and at the 2.5-hour timepoint in **Figure 4B** Spike still had rounded plus strand only condensates as indicated by yellow arrows. We thus reasoned that there must be some difference in the affinity of plus and minus strand RNA duplexes for Spike as compared to the others and Spike might also be the least “sticky” duplex. Of note, we previously observed that SARS-CoV-2 had higher GC content at viral ends (e.g. Nucleocapsid RNA) then middles (e.g. Spike RNA)^21^. This arrangement was not consistently observed in other viruses regardless of type (**Supplemental Figure S4E**). Thus, our results suggested that a comparison of Spike RNA to other tested RNA features may provide insight into the sequence specificity controlling trans RNA-RNA interaction.

To assess if Spike plus/minus RNA is different from other tested RNA duplexes we began with an examination of the primary sequence content. We observed that of our tested TRS-B fragments, Spike had the lowest GC content (**Figure 4F**) and further, the degree of predicted - MFE energy of hybridization^45^ was perfectly correlated with the GC content suggesting this is the least sticky RNA sequence (**Figure 4G**). Similar results were also observed in RNA only simulations in **Supplemental Figure S3**.

We reasoned if depletion in GC content was an RNA viral strategy for reducing RNA–RNA interactions then depletion in GC content should be a near universal feature of RNA viral genomes. To this end, we computed the GC content by RNA virus type and, consistent with previous publications^61^, we observed that RNA viruses tended to have lower GC content than that of the human transcriptome (**Figure 4H**). Intriguingly, the RNA virus type with the lowest GC content was negative strand RNA virus which must first replicate a plus strand copy prior to protein production. Negative strand viruses must arguably have the greatest problem with stickiness as no translation occurs prior to plus strand genome synthesis^62^ event to potentially prevent RNA– RNA interaction. We examined the patterning of the GC content to see how common GC content depletion was in RNA viruses. Intriguingly, only in a few viral orders including nidoviruses was the GC content in the at the end of the plus strand greater than that at the middle; the majority of viruses displayed the opposite configuration (**Supplemental Figure S4E**). These data collectively suggest that nidovirus genomes are particularly depleted in GC content at the middle and that this may help mitigate RNA–RNA interaction between the plus and minus strand genomes during replication.

Our data from **Figure 3 and 4** suggest that RNA viral sequences may be under selective pressure to maintain low GC content to mitigate RNA–RNA interactions between plus and minus strand genomes during replication. Thus, we reasoned that as viruses may be under selective pressure to maintain low GC content this should lead to a commensurate decrease in the propensity to form g-quadruplexes. To examine this possibility further we predicted g- quadruplexes sequences^63^ in the SARS-CoV-2 genome. In keeping with previous observations of acute RNA viruses^64^, we observed that SARS-CoV-2 is overall has both few and low-quality predicted g-quadruplex sequences^65^ (**Supplemental Figure S4F, G and H**) with no predicted sequence remotely approximating the score of a bona fide RNA g-quadruplex, Orn-1^66^. Additionally, only one tested RNA fragment, the 5′end plus strand, had a predicted g-quadruplex sequence (**Supplemental Figure S4G and S4H**). Addition of non-physiological excessive^66^ magnesium ion concentrations (10mM) had no significant impact on the degree of co-localization under conditions of staggered minus strand RNA addition. (**Supplemental Figure S4I and S4J**). Thus, we concluded that g-quadruplexes do not contribute significantly either to the arrested morphology of the condensates or the degree of RNA–RNA interactions.

### Control of RNA-RNA interactions by altering the ratio of plus and minus strand RNA

Although low-GC content was correlated with a reduction in mixing of RNAs and a reduction in RNA–RNA interaction strength, low GC content and N protein coating are in combination still insufficient to block the formation of arrested networks (**Figure 3 and 4**). We wondered if there might be other strategies employed by RNA viruses to limit the spurious interaction between the plus and minus strand RNA genomes during viral replication. Plus-strand RNA viruses like SARS-CoV-2 begin with a single complete plus strand genome carried by the virion^48^. From this plus strand RNA, at least one minus strand template is produced^67^.

Therefore, at early times in infection, plus and minus RNA are perhaps present at equal molar ratios, however this ratio does not persist over the duration of infection as the virus uses the minus strand RNA to produce numerous copies of plus strand RNA. Thus, as infection progresses, a higher proportion of plus strand is produced relative to minus strand. We hypothesized if the change in this ratio might also mitigate the strong RNA–RNA interactions by effectively spreading plus/minus RNA–RNA interactions over an increasing higher number of plus strand molecules such that eventually the contributions of the minus strand become effectively saturated.

To test the hypothesis that the ratio of plus/minus RNA strands can influence the morphology and arrest of condensates, we produced condensates with 1:1 plus/minus, 10:1 plus/minus and 100:1 plus/minus RNAs. Of note, in all cases 100nM of total RNA and 3.2μM of protein were used. We observed that in all tested TRS-B fragments, higher ratios of plus strand RNA relative to minus resulted in a suppression of the arrested network formation with a 10:1 partially and 100:1 almost completely suppressing the amorphous shape/ lack of circularity observed when 1:1 ratio are present (**Figure 5A**).

**Figure 5.**
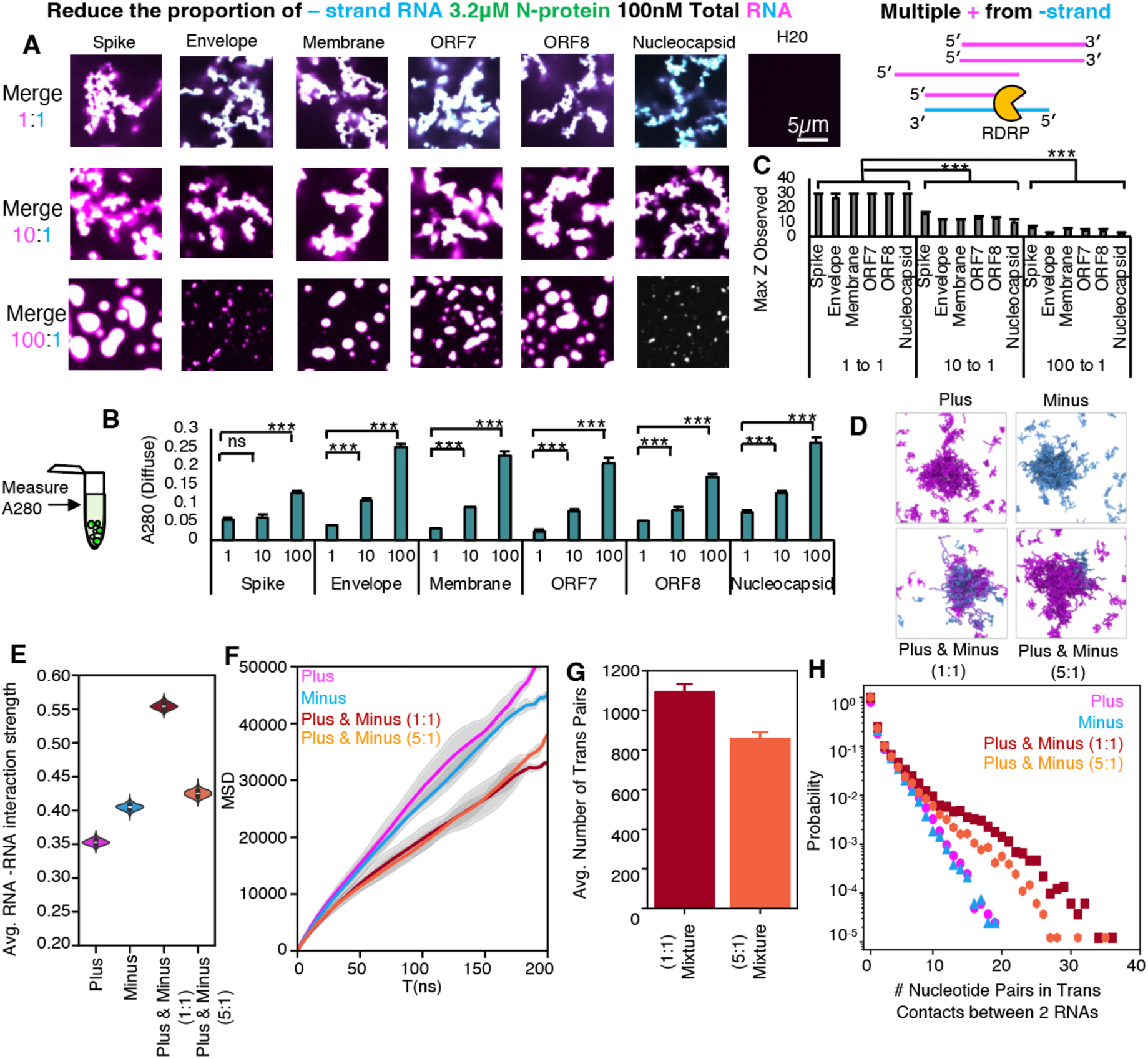
Imbalance in complementary RNA sequence concentrations present in late-stage viral replication attenuates RNA-RNA interactions **(A**) Cartoon depicts the RDRP (orange Pacman) synthesizing multiple plus strand RNA genome copies from a single minus strand RNA indicating plus and minus strand RNA genome ratios are not uniform over the course of infection. 1:1 ratio mix (early infection) of plus and minus strand RNA result in arrested networks and 100 plus (magenta) to minus (cyan), consistent with later in infection, suppress arrested network formation resulting in more liquid-like condensates. 10:1 plus to minus result in intermediate phenotypes. **(B)** Diffuse phase RNA measurements related to panel A. Higher plus strand to minus strand RNA ratio results in a commensurate increase in RNA and protein signal (A280) absorbance in the diffuse phase indicating the lower ratios are better able to drive condensation. Error bars are technical replicates. NS not significant (p<0.001 ***) **(C)** Related to **Supplemental Figure S5**. Observed Z height for condensates formed in A. 1:1 ratios of plus and minus RNA extend at least 30 microns into Z (arrested networks) whereas 100:1 extend only a few microns. 10:1 is intermediate. Error bars are 3 technical replicates. (p<0.001 ***). **(D)** Comparison of 2X plus, 2X minus, 1:1 plus/minus, and 5:1 plus minus in RNA only simulations of envelope RNA (model 2). **(E)** Average RNA-RNA interaction strength for systems depicted in (**D**). Consistent with experiments, 1:1 plus/minus (maroon) results in a stronger interaction (i.e., most negative) than the 5:1 (orange) system. **(F)** 5:1 plus/minus simulated RNA system (orange) shows intermediate MSD between 2X plus (magenta) or 2X minus (cyan) and 1:1 plus/minus (maroon). **(G)** Reduction in trans RNA–RNA pairs between 5:1 plus/minus and 1:1 plus/minus in RNA simulations. **(H)** Probability of long trans RNA-RNA interactions decreases in 5:1 as compared to 1:1.

Consistent with the morphology change, the plus/minus RNA ratio alters the amount of RNA in the dilute phase (**Figure 5B**), with excess plus strand associated with more RNA in dilute phase, despite total RNA in the reactions being the same. To quantify the degree of network formation, we measured the height of a continuous condensate from the cover glass up into solution. We found in 1:1 mixes that the arrested network extends from the glass up to 30 microns into Z **(Figure 5C** **and Supplemental Figure S5A, S5B, and S5C**) with 10:1 extending an intermediate height into solution as compared to 1:1 and 100:1 indicating a suppression of arrested network formation with increasing plus/minus RNA ratios. We were curious as to how changing RNA ratio altered the arrangement of molecules in the dense phase. To characterize the impact of the mixing ratio on the strength of RNA–RNA interactions and RNA strand dynamics, at the nucleotide level, we simulated a system composed of a 5:1 plus/minus strand ratio (via model 2, see Methods, **Figure 5D**). Indeed, an excess of plus strands yields reduced RNA–RNA interaction strength (**Figure 5E**) and exhibits slightly faster dynamics on long timescales (**Figure 5F)** compared to the 1:1 plus/minus case. These effects can be attributed to a significant reduction in trans RNA contacts when plus strands are in excess (**Figure 5G and H**). These data collectively indicate that the degree of RNA–RNA interactions contribute to arrested network formation and that altering the ratio of plus and minus RNA may be an additional strategy used by RNA viruses to mitigate complementarity-induced arrested network formation.

Collectively, our data thus far suggest that perfect complementarity between the plus and the minus strand leads to arrested network formation proportional to the RNA–RNA interaction strength as governed by N protein interactions (**Figure 4**), RNA sequence and in particular GC content (**Figure 4**) and the ratio of RNAs present (**Figure 5**). These combined features serve to minimize entanglements by reducing hydrogen bonding between complementary RNAs and spreading of RNA interactions over increased number of plus strand RNA molecules relative to minus. We wondered if other strategies might be employed to reduce complementarity perhaps via a change in RNA sequence for one of the pairs of RNA as each of the above strategies was not completely sufficient for preventing arrested condensate morphology.

### Reducing RNA-RNA interaction strength with mutations or staggering RNA overlap reduces arrested network formation

As RNA viruses replicate, they accumulate mutations in their genomes due to the high error rate of viral RNA dependent RNA polymerases (e.g. average of 2.68 × 10^−5^ de novo errors per cycle with a C > T bias for SARS-CoV-2 RdRp^68^) and the action of RNA modifications such as inosine (ADAR^69^), methylation (METTL3/14)^69^, and 2’OME^70^ (NSP16)^71^. Notably, all these modifications have been detected on SARS-CoV-2 viral RNA. These mutations/modifications may offer a selective advantage to RNA viruses by increasing the diversity of the genome allowing for escape from immune surveillance. In keeping with this theory, most recent pandemics were caused by RNA viruses^72^ with the single most recent DNA virus pandemic, Mpox, arising due to reduced frequency of individuals vaccinated for smallpox in the population^73^. Thus, increasing mutational burden represents a positive effect for RNA viruses with a tradeoff between increased diversity and continued functionality of protein and RNA elements but potentially a secondary benefit of reducing complementarity between the plus and minus strands of the RNA genome.

Since RNA viruses accumulate multiple mutations/modifications during replication, we wondered if these changes might also be a strategy employed by RNA viruses to reduce RNA– RNA interactions between the plus and the minus strand RNAs. Thus, we sought to ask, can mutations on minus RNA strand reduce complementarity to the plus RNA strand and if so, how many mutations are required?

To this end, we chose to reduce complementarity to the plus strand through modifications of minus strand RNA in two ways, **(1)** increased inosine content and **(2)** increased mutation in the template sequence. We chose the minus strand rather than the plus as this RNA had less propensity to drive condensation with N protein on its own (**Figure 1**).

We observed that increasing amounts of inosine (**Figure 6A**) did little to mitigate the arrested network morphology of the condensates but, following 4-5 rounds of error prone PCR, arrested network formation was partially mitigated (**Figure 6B**). Of note, moving forward we limited our experiments to two RNA pairs as Spike and Nucleocapsid RNA TRS-Bs represent the least and most “sticky” RNA combinations, respectively as measured by GC content. (**Figure 4F**) Thus, these results should be representative of the possible RNA conditions for any perfectly complementary RNA pair.

**Figure 6.**
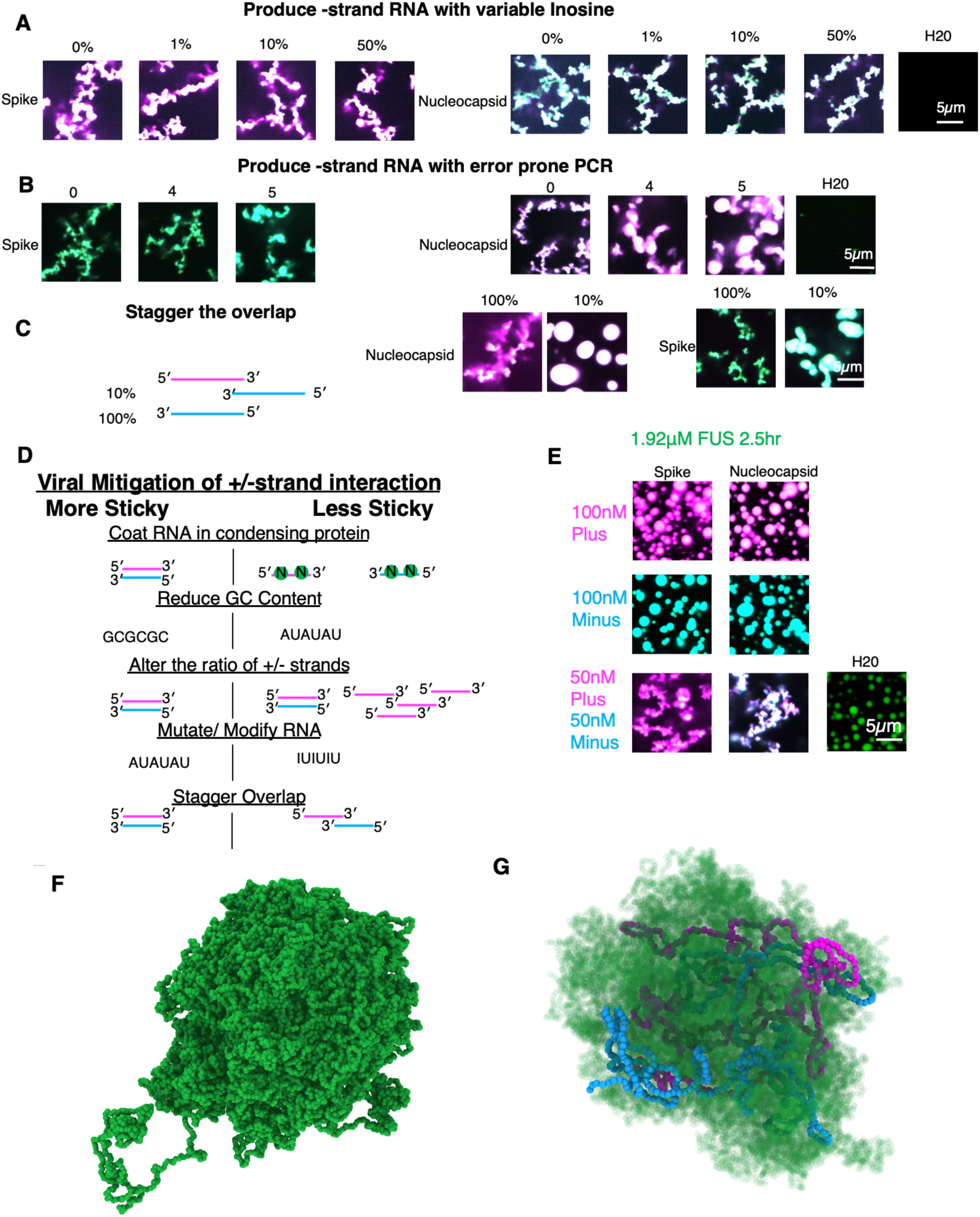
Protein sequence independent role for RNA–RNA interactions in driving biomolecular condensate morphology. **(A)** No obvious effect of increasing inosine concentration on arrested network morphology. Minus strand RNA (cyan) was produced with 0, 1, 10, and 50% inosine to replace increasing amounts of A nucleotides with I. Combinations of two RNAs are 50nM each. Scale bar (white line in H_2_O water only control) is 5 microns. All images in each panel are acquired on the same day and are contrasted the same. **(B)** 4-5 rounds of error prone PCR are sufficient to reduce the arrested network formation. Minus strand RNA (cyan) was produced with DNA templates made from 0 (I proof) or multiple rounds of error prone PCR. 1μL of error prone PCR reaction one was used as template for error prone PCR 2. Of note cycles 1-3 were also tested and resulted in no obvious change in arrested network formation. **(C)** Reducing the overlap of the plus and minus RNAs by shifting the position of the minus strand RNA template results in condensates with more circular morphology. **(D)** Summary of multiple viral strategies used to avoid extensive RNA–RNA interactions. **(E)** FUS protein also yields arrested networks when perfectly complementary plus and minus strand RNA is present. 100nM of plus (magenta, top row) and minus (cyan, middle row) strand RNA from Spike and Nucleocapsid TRS-Bs yielded rounded co-condensates with FUS protein (green). 50nM each plus (magenta) and minus (cyan) yield arrested networks (bottom panel) with FUS. All images depict the merged signal of the same total RNA (100nM) and N protein (3.2μM) concentrations. Scale bar (white line in H2O water only control) is 5 microns. All images in each panel are acquired on the same day and are contrasted the same. Simulations of FUS and RNA. **(F)** Simulation of full 64xFUS proteins without RNA. The proteins can form condensates. **(G)** Simulations of 64xFUS proteins (semitransparent green) with two Envelope RNA plus (magenta) and minus (blue) strands. The FUS proteins can colocalize with the protein.

Our data thus far revealed that the presence of perfectly complementary stretches of 480 nucleotides results in arrested network configurations in condensates. As such long stretches of complementarity are probably actively avoided by host cells by careful organization of the genome (e.g., splicing, non-overlapping genes in Watson and Crick DNA sequences, staggered expression timing, RNA export etc.),we reasoned it would be informative to reduce the complementarity by staggering the overlap of the sequence to be 10 percent of the previous conditions (48 nucleotides of perfect complementarity sequences). We did this by altering the positioning of the template for the minus strand in the genome shifting it 3 prime to the plus strand RNA template. We observed that the 10 percent overlap condition almost completely suppressed the arrested network phenotype, suggesting once again that the degree of RNA– RNA interaction strength is proportional to the arrested morphology (**Figure 6C**). In keeping with this finding, we would expect that producing sub-genomic RNA fragments^34^ (sgRNA), a strategy employed by betacoronavirus might be yet another strategy to reduce interaction between plus and minus strand RNA sequences as these shorter sequences would have less propensity to base pair than full-length genomes. Thus, multiple strategies to mitigate RNA–RNA interactions between perfectly complementary RNA sequences are accessible to viruses (**Figure 6D**).

### Protein content is dispensable for morphology regulation by extensive RNA-RNA interaction

Given that 48 nucleotides had only modest degree of conglomeration, we reasoned that in any given eukaryotic cell, perfect sense and antisense RNA should only be problematic if the number of identical nucleotides is greater than 48 nucleotides, particularly when both sequences are simultaneously transcribed in the same nucleus. Viruses replicate not only in the presence of their own proteins but in the milieu of many host factors including an array of host- RBPs.

As Fused in sarcoma (FUS) protein is the best characterized condensing protein which can bind RNA sequences with high nuclear protein concentrations^74^, we reasoned this would be an ideal candidate to test whether strong RNA–RNA interactions drive arrested network formation regardless of the identity of the co-condensing protein. To this end we tested our two representative RNA duplexes plus/minus Spike and Nucleocapsid RNA TRS-B and plus/minus with FUS protein rather than N protein and observed that, in keeping with the results for N protein, FUS was able to make round condensates with either plus or minus Spike or Nucleocapsid RNA but formed arrested networks with the combination of the plus and minus RNA (**Figure 6E**). These data suggest that RNA–RNA interaction network duplex formation is not dependent on condensing protein identity and thus should be a universal feature of biomolecular condensates. These data suggest condensate morphology is due to RNA–RNA rather than RNA–protein or RNA–N protein interactions. Simulations of FUS protein without (**Figure 6F**) or with RNA (**Figure 6G**) further illustrated that FUS protein could reduce RNA– RNA interactions when complementary sequences were present.

### Role of cytoplasmic proteins in modulating RNA-RNA interactions

Lastly, we wanted to know if the observation we saw cell free also held true in cells which contain many other host proteins (including FUS) and RNAs. To this end, we introduced N protein and our RNA duplexes into the cytoplasm of Vero E6 cells utilizing acid washed beads^75^ to deform the membrane allowing for the penetration of RNA and protein (**Figure 7A**).

**Figure 7.**
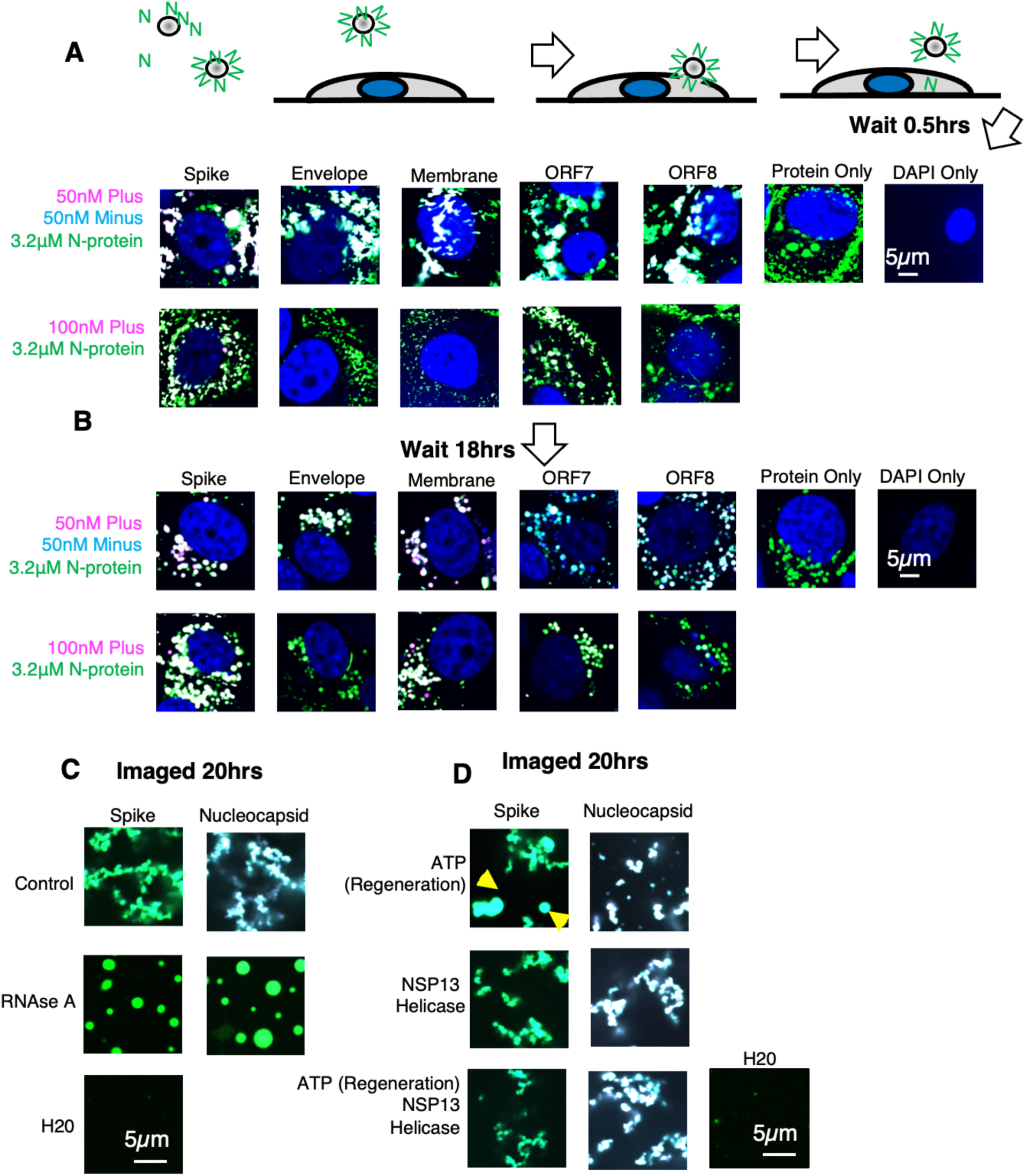
Cytoplasmic RNAses and Helicases may help to resolve arrested networks. **(A**) Cartoon depicts acid washed bead-based strategy for the delivery of RNA and protein to the cytoplasm of cultured, adherent, mammalian cells. Beads are incubated with protein (green Ns) and RNA solution in physiological buffer. They are then rolled over the membrane of adherent mammalian cells. Contact between the bead and the membrane surface disrupts the membrane such that protein and RNA can enter. Resulting protein is deposited in the cytoplasm (grey semi-circle) not the nucleus (blue circle). Representative images showing the merged signal (plus RNA in magenta, minus RNA in cyan, N protein in green, nucleus in blue). From 5 tested RNA duplexes (top panel) or just plus strand RNA (bottom panel). Imaged at 0.5 hours after bead treatment. All duplexes tested result in arrested networks (top panel) whereas plus alone does not (bottom panel). **(B)** Representative images showing the merged signal (plus RNA in magenta, minus RNA in cyan, N protein in green, nucleus in blue). From 5 tested RNA duplexes (top panel) or just plus strand RNA (bottom panel). Imaged at 18.5 hours after bead treatment. Compared to the 0.5-hour timepoint (A) condensates appear to relax. **(C)** Treatment with RNAse A resolves arrested networks. Representative images showing the merged signal (plus RNA in magenta, minus RNA in cyan, N protein in green), **(D)** Presence of ATP in the absence of SARS-CoV-2 RNA helicase NSP13 results in partial mitigation of arrested network formation for spike (yellow arrows).

We observed that 1 hour following incubation with beads, protein and RNA, all tested RNA duplexes had observable arrested networks formed in the cytoplasm of cells containing both protein and RNA signals. Of note, we did not test Nucleocapsid RNA to avoid any potential contributions from the additional protein translation (our protein was produced with the native SARS-CoV-2 sequence and therefore may contain small amounts of contaminating plus RNA sequence from bacterial production of Nucleocapsid RNA and in cells this could be translated to additional N protein fragments). In comparison, addition of only plus strand RNA and N protein resulted in more rounded condensates. The cytoplasm of eukaryotic cells contains numerous RNA transcripts, ribosomes, and proteins (RNA-binding or otherwise). To address if incubation in the cytoplasmic milieu could resolve the arrested network we again imaged cells 18 hours later (**Figure 7B**). We observed that, following 18-hour incubation, N protein and RNA signal no longer resembled an arrested network but rather more rounded condensates. Similar rounded condensates were observed in plus strand RNA only conditions. Collectively these data suggested that one or more cellular factors may be acting on the RNA and protein arrested networks at long time scales to resolve them.

To address what mechanisms could be used by the cell to resolve an arrested network of RNA–RNA interactions we returned to our cell free system. As double stranded RNA present in our RNA duplexes of a viral origin should be an excellent target for the host native immune system^76^, we first reasoned that one pathway for resolving arrested networks may be to simply degrade the RNA through the action of endogenous RNAases such as the interferon induced RNAse L etc^77^. In SARS-CoV-2 infections, this activity could be provided by NSP15 which preferentially degrades AU rich dsRNA^78^. To test this possibility, we added RNAse A to our preincubated duplexes and N protein (0.5 hours) (**Figure 7C**). We observed that addition of RNAse A was sufficient to completely abolish the arrested network phenotype, indicating that an additional way that cells may resolve an arrested network is to degrade the RNA holding it together via its extensive network of RNA–RNA interactions.

We reasoned another likely candidate protein family that could act to resolve the arrested networks was “helicases” which are thought to function by unwinding RNA duplexes in some cases^79^. Contained within the genome of SARS-CoV-2 is the RNA helicase NSP13^80^. This protein is essential for viral RNA replication. Although not present in uninfected Vero cells, we reasoned that this protein was the most likely candidate for resolving SARS-CoV-2 RNA–RNA interactions during infection. To test if NSP13 could resolve an RNA duplex of 480 nucleotides long (far longer than any duplex tested for any helicase activity to our knowledge), we incubated our Spike and Nucleocapsid TRS-B plus/minus RNA duplexes with N protein and NSP13. As NSP13 activity is dependent on ATP and ATP is a reported hydrotrope^81^ which can promote solubilization of biomolecular condensates we also included +/- ATP/creatine kinase ATP regeneration as controls (**Figure 7D**). Intriguingly, NSP13 alone had no noticeable effect on the arrested network formation nor did NSP13 with the addition of ATP. However, the addition of ATP alone did have a modest effect on the morphology of Spike TRS-B plus/minus arrested networks in keeping with the published observation of ATP’s hydrotrope activity. This effect of ATP was not readily observed in Nucleocapsid TRS-B RNA conditions consistent with Spike RNA being the “least sticky” with respect to GC content (**Figure 4F**). We interpret these results as indicating that ATP’s hydrotrope activity can help limit weaker RNA–RNA arrested network formation only if it has not been consumed by NSP13 which has reported tremendous ATPase activity^82^. Thus, high intracellular ATP may help solubilize arrested networks but once a 480- nucleotide duplex is formed it cannot be resolved under tested conditions by NSP13.

## DISCUSSION

Beyond storing information, RNA provides an essential architectural role in the formation of biomolecular condensations^24^. Viruses risk RNA-RNA interactions that could lead to potentially deleterious states, these are the strategies we have uncovered that viruses can use to avoid the risks inherit in complementary RNA by (1) pre-coating RNA genomes in condensing protein, (2) reducing genome GC content, (3) altering the ratio of plus and minus strands, and (4) Mutating or modifying RNA. This concept likely extends to all eukaryotic condensates which can additionally (5) tune native RNA-RNA interactions in condensates by stagging overlapping gene sequences in DNA. Thus, in our manuscript, we present a likely universal feature of RNA containing biomolecular condensates, uncovering how RNA–RNA interaction complementarity confers the morphology of the resulting condensate. We uncovered this central axiom of biomolecular condensates by reconstituting representative molecules from two essential SARS-CoV-2 replication steps: subgenomic RNA generation and genome replication. We observed that low affinity interactions which arise because of low sequence complementarity (e.g. pairing between the plus TRS-L and minus TRS-B) yield rounded condensates whereas high affinity interactions (e.g. pairing between the plus TRS-B and minus TRS-B) due to longer, stronger complementarity RNA sequences yield arrested networks. Condensate morphology correlates with experimental measurements of RNA–RNA interaction strength via direct sequencing of reduced DEPC reactivity and via simulations.

We believe this feature of RNA and condensate biology was not yet discovered due to (1) the overall reliance on the field on shorter and homotypic RNA polymers (2) the preference for biochemical reconstitution experiments to be undertaken with a single RNA species for ease of interpretation. These previously implemented conditions would not be predicted to yield arrested networks unless RNA structure is extraordinarily low^37,38^ or multiple RNAs are present as is the case for reconstituted stress granules^83,84^ from cellular RNA extract. Our work provides a unique insight into this RNA dependent process as it explores the RNA parameters required to yield entangled RNA. RNA–RNA interactions are present in all condensates but, according to our model interesting changes to condensate morphology happen when a high degree of complementarity is present. This in principle can occur when 2 or more RNAs are present (as with our experiments in this paper) or when longer sequences of RNA are present (the longer the RNA the more chances for complementarity). Our experimental and modeling data suggest that arrested network morphology is due to contributions from trans RNA contacts and long runs of RNA contact rather than protein–RNA or protein–protein interactions or random RNA entanglement making this a universal feature of RNA containing biomolecular condensates. Our simulation data suggests that these networks arise predominately from the daisy-chaining of incomplete base pairing between the complementary RNA strands, favoring arrested networks over perfect duplexes. It is likely that both RNA structure and protein binding also serve to coat regions of RNA blocking perfect complementarity to promote arrested network formation.

Our results suggest that lowered GC content confers reduced RNA–RNA interaction strength. RNA viral genomes are dramatically depleted in GC content. Of note, the most GC depleted order of plus strand RNA viruses following nidovirales like SARS-CoV-2 is Cryppavirales. This is intriguing as many constituent species from Cryppavirales have lost all packaging machinery (e.g. capsid/nucleocapsid proteins)^85^. These viruses may instead rely solely on reduced GC content to prevent sticking during replication. Further evidence of the selective advantage of low GC content has been observed in RNA sequencing data collected over the course of the pandemic. These data reveal that SARS-CoV-2 has accumulated numerous coding and non-coding mutations over time. Intriguingly, the virus has undergone excessive C–U transitions^86^, likely mediated by host-cell RNA editing machinery, specifically mediated by APOBEC proteins^87^.

Our results suggest that SARS-CoV-2 and other viruses may benefit from the lower GC content during replication with edited genomes providing a selective advantage to the viruses during replication beyond the increase in genetic diversity by reducing extensive RNA–RNA interactions. In keeping with this observation, we observed that addition of inosine, the product of ADAR mediated editing) to minus RNA had limited impact on the propensity to form arrested networks. We would speculate this is due several factors; **(1)** inosine chemically resembles guanosine which can pair with both C and U^88^, **(2)** the addition of inosine could disrupt minus strand RNA structure, rendering RNA more single stranded and prone to base pairing with plus, **(3)** error prone PCR results suggest that more than 60 mutations or 12.5 percent (50 percent of average A content) of nucleotides need to be mutated before there is a noticeable impact on A content. Of note, for our error prone PCR conditions, 5 cycles should result in approximately 200 mutations per RNA molecule, not confined to A nucleotides. This may explain the observed net loss of C bases^86^ as opposed to A bases as our results suggest that ADAR editing may increase RNA–RNA interactions strength whereas APOBEC mediated C–U transitions may act to reduce RNA-RNA interaction strength.

Viruses including coronaviruses actively maintain the structure of the RNA in defined configurations when mutations arise through compensatory mutations to restore base pairing (such as the replacement of a GC pairs with an AU or GU wobble)^89^. Viruses thus retain RNA structure despite it being a trigger for innate immune surveillance. It is intriguing to speculate that viruses employ RNA structure as one part of their multi-pronged strategy of mitigating extensive RNA–RNA interactions, burying the complementary sequence in cis RNA structure to avoid pairing in trans.

Other parts of the viral strategy to mitigate strong RNA–RNA interactions involve coating RNA in condensing protein and altering the ratio of plus and minus strands and we have presented evidence for both in reducing arrested network formation. We postulate that these strategies are also likely applicable to non-plus stranded RNA viruses. Our results suggest that the replication strategy employed by all RNA viruses to produce proportionally fewer template RNA molecules relative to packaged RNAs may have an additional advantage beyond economy, as this strategy limits the damaging effects of extensive RNA–RNA interactions, spreading perfect complementarity over an increasingly large number of molecules. Additionally, our results suggest that a reason for the enrichment of proteins predicted to be prone to condensation in viral genomes^90,91^, may be to prevent extensive RNA–RNA interactions during genome replication by separately coating the genome and its template. Minus strand RNA viruses may rely more heavily on genome condensation and condensing proteins to block extensive RNA-RNA interactions whereas plus and double stranded RNA viruses may employ ribosome mediated translation to separate their genomes. This may explain why minus strand RNA viruses are more likely to employ inclusion bodies (a biomolecular condensate formed by the virus during genome production) in their replication cycle^92^.

Our helicase experiments suggest that preformed genome-long duplexes may be difficult to resolve by even viral machinery. How then can the replication of double-stranded RNA viruses be explained, which contain perfect genome-long double-stranded RNA duplexes packaged in the virion? We believe the answer to this may be found in the packaging and genome organization strategies employed by double-stranded RNA viruses. Double stranded RNA virus genome length tends to be shorter and split over multiple molecules, both features which we would predict would reduce RNA-RNA interactions. As to packaging, dsRNA utilizes a unique packaging strategy as compared to plus and minus strand RNA viruses. Chronologically, plus strand RNA viruses are thought to have evolved first, followed by double-stranded RNA viruses which then give rise to minus strand RNA viruses^93^. In keeping with this model, double-stranded RNA viruses are initially packaged in a manner like plus strand RNA viruses but following packaging of single copies of plus strand RNA, complementary minus strand RNA is replicated in the virion by pre-loaded RDRP^94^. Thus, +/- strand genome entanglement may be limited by reducing the number of molecules, restricting minus strand synthesis to the virion, and by near simultaneous synthesis of all RNA duplexes.

In cells and during infection helicases/RNAses as well as endogenous ATP may help to resolve duplexes. We will briefly speculate on the implications of these findings in the following sections. <u>RNAses</u>: We would speculate that endogenous and virally encoded (e.g. NSP15 in SARS-CoV-2^78^) RNAses may be a strategy of last resort to resolve RNA tangles with the negative consequence of degradation of RNA. In keeping with this theory, NSP15 is an essential protein for SARS-CoV-2 viral replication, and RNAseL mediated degradation can dissolve most RNA-dependent condensates in cells^77^. <u>Helicases</u>: Under our tested conditions, NSP13 was unable to resolve the arrested network of RNA. We do not preclude the possibility that under some conditions NSP13 could prevent or reduce extensive RNA-RNA interaction as suggested by its essential role in viral replication, rather our data indicate the possibility that there are some conditions which might be impossible to resolve. It is also possible that in our reconstitution experiments NSP13 is missing one or more required co-factor required for rapid processivity.

Eukaryotic cells may partially avoid the problem of long RNA-RNA interactions in native RNA sequences by organizing their genomes in such a way as to avoid perfectly complementary stretches of RNA, but a nuclear protein could further help to limit RNA–RNA interaction. We postulated that a high-concentration, nuclear-localized, condensing RNA-binding protein might perform a similar role endogenously in eukaryotic transcription as SARS-CoV-2 N protein does for viral RNA production. Following this logic, we reasoned that host condensate proteins may also result in arrested networks upon binding to perfectly complementary RNA. Thus, we sought to test the propensity of such a protein to form arrested networks when perfectly complementary RNA is present and show that FUS may be partially able to mitigate extensive RNA-RNA interaction in a manner like N. These data collectively suggest a protein sequence independent role for condensing proteins in suppression of strong trans RNA-RNA interactions.

In summation, our data collectively reveal the importance of RNA-RNA interaction strength in regulating condensate morphology. Weak RNA-RNA interactions conferred by low complementarity in RNA sequences results in round condensates whereas strong RNA-RNA interactions conferred by high complementarity in RNA sequences resulted in arrested networks. Our data indicate this phenomenon is largely RNA and protein sequence independent and thus broadly applicable across the biomolecular condensate field whenever RNA is present. Additionally, our results suggest explanations for the enrichment for specific RNA sequence and structure features in RNA viral genomes due to reliance on RdRp proteins and RNA templates for replication.

## ACKNOWLEDGEMENTS

We thank Professor Yifan Dai for the use of his microscope and his lab in the collecting the preliminary data for this manuscript. We thank Dr. Wilton Snead for providing FUS and TEV protein. We thank Professor Zhao Zhang in the Department of Pharmacology and Cancer Biology for use of his Nanopore Instrumentation GridIon. This study was supported in part by the Center for Microbial Pathogenesis and Host Inflammatory Responses grant P20GM103625 through the NIH National Institute of General Medical Sciences Centers of Biomedical Research Excellence.

J.C.M and K.D.R were supported by NIGMS MIRA (R35GM122601) and a kind gift from the Palade Family. J.A.J. acknowledges start-up funds provided by the Department of Chemical and Biological Engineering and the Omenn–Darling Bioengineering Institute at Princeton University. J.A.J. also acknowledges research support from the Chan Zuckerberg Initiative DAF (an advised fund of Silicon Valley Community Foundation; grant 2023-332391), the National Institute of General Medical Sciences of the National Institutes of Health under Award Number R35GM155259, and the National Science Foundation (NSF) through the Princeton University (PCCM) Materials Research Science and Engineering Center DMR-2011750. The simulations reported on in this manuscript was substantially performed using the Princeton Research Computing resources at Princeton University, which is a consortium of groups led by the Princeton Institute for Computational Science and Engineering (PICSciE) and Office of Information Technology. C.A.R .acknowledges start-up funds provided by the Université de Montréal. C.A.R. was additionally supported by NIH T32CA9156-43, F32GM136164, K99AI173439-01A1 and L’OREAL USA for Women in Science Fellowship. V.Z. and A.S.G were supported by 7R01GM081506-13. V.Z. was supported by Duke University School of Medicine International Chancellor’s scholarship.

## COMPETEING INTERESTS

All other authors declare that they have no competing interests.

## DATA AND MATERIALS AVAILABILITY

All data and/or materials are available upon request from C.A.R. or J.A.J.

## CONTRIBUTIONS

J.A.J, C.A.R., and A.S.G. conceptualized the project, designed experiments, prepared figures, drafted and edited the manuscript. C.A.R. and V.Z. performed experiments, prepared figures and analyzed data. D.A. and R.S. designed and performed experiments and computational analyses, analyzed data, prepared figures. J.C.M and K.D.R designed experiments and analyzed data. All authors contributed to the editing of the manuscript.

## METHODS

Protein production: SARS-CoV-2 nucleocapsid protein was produced occurring to our established protocols^21^. RNAse A was purchased from NEB (T3018L).

**FUS protein production and labeling** We produced the following sequence. **6xHis-MBP-FUS FL WT.**

MGSDKIHHHHHHSSGTKIEEGKLVIWINGDKGYNGLAEVGKKFEKDTGIKVTVEHPDKLEEKFPQVAATGDGPDIIFWAHDRFGGYAQSGLLAEITPDKAFQDKLYPFTWDAVRYNGKLIAYPIAVEA LSLIYNKDLLPNPPKTWEEIPALDKELKAKGKSALMFNLQEPYFTWPLIAADGGYAFKYENGKYDIKDVGVDNAGAKAGLTFLVDLIKNKHMNADTDYSIAEAAFNKGETAMTINGPWAWSNIDTSKVNYGVTVLPTFKGQPSKPFVGVLSAGINAASPNKELAKEFLENYLLTDEGLEAVNKDKPLGAVALKSYEEELAKDPRIAATMENAQKGEIMPNIPQMSAFWYAVRTAVINAASGRQTVDEALKDAQTNSGSDITSLYKKAEGGTENLYFQGHMASNDYTQQATQSYGAYPTQPGQGYSQQSSQPYGQQSYSGYSQSTDTSGYGQSSYSSYGQSQNTGYGTQSTPQGYGSTGGYGSSQSSQSSYGQQSSYPGYGQQPAPSSTSGSYGSSSQSSSYGQPQSGSYSQQPSYGGQQQSYGQQQSYNPPQGYGQQNQYNSSSGGGGGGGGGGNYGQDQSSMSSGGGSGGGYGNQDQSGGGGSGGYGQQDRGGRGRGGSGGGGGGGGGGYNRSSGGYEPRGRGGGRGGRGGMGGSDRGGFNKFGGPRDQGSRHDSEQDNSDNNTIFVQGLGENVTIESVADYFKQIGIIKTNKKTGQPMINLYTDRETGKLKGEATVSFDDPPSAKAAIDWFDGKEFSGNPIKVSFATRRADFNRGGGNGRGGRGRGGPMGRGGYGGGGSGGGGRGGFPSGGGGGGGQQRAGDWKCPNPTCENMNFSWRNECNQCKAPKPDGPGGGPGGSHMGGNYGDDRRGGRGGYDRGGYRGRGGDRGGFRGGRGGGDRGGFGPGKMDSRGEHRQDRRERPY

The 6xHis-MBP-FUS FL WT (MBP-FUS) and MBP-7xHis-TEV protease (MBP-TEV) plasmids were obtained from Addgene (plasmids # 98651 and # 8827, respectively). After transforming the plasmids into BL21 E. coli, cells were grown at 37°C until reaching OD600 0.6-0.8 and proteins were induced overnight at 18°C following addition of IPTG (0.5 mM and 0.4 mM for MBP-FUS and MBP-TEV, respectively). Cells were harvested by centrifugation at 14,000 rcf at 4degC the following day and resuspended in lysis buffer. For MBP-FUS, lysis buffer consisted of 50 mM HEPES pH 7.4, 1.5 M NaCl, 10 percent glycerol, 20 mM imidazole, 5 mM beta-mercaptoethanol, and EDTA-free protease inhibitor cocktail (Pierce A32965); for MBP-TEV, lysis buffer consisted of 25 mM HEPES pH 7.4, 300 mM KCl, 10 percent glycerol, and 20 mM imidazole. Cells were lysed using probe sonication (QSonica Q500 with 20 kHz converter and 1/4” microtip), and lysate was clarified by centrifugation at 27,000 rcf for 30 min at 4°C. Clarified lysate was mixed with 0.4-0.5 mL of washed, packed HisPur cobalt resin (Thermo Scientific 89965) per 1 L of cells for 1 hour at 4°C. After protein binding, resin was transferred into a gravity flow column and washed with approximately 80-100 resin bed volumes of lysis buffer without protease inhibitor cocktail. MBP-TEV was further washed using approximately 100 resin bed volumes of lysis buffer containing 10 mM ATP-MgCl2, followed by an additional 80-100 resin bed volumes of lysis buffer without ATP-MgCl2. Proteins were then eluted using elution buffer. For MBP-FUS, elution buffer consisted of 50 mM HEPES pH 7.4, 150 mM NaCl, 10 percent glycerol, 200 mM imidazole, and 5 mM beta-mercaptoethanol; for MBP-TEV, elution buffer consisted of 25 mM HEPES pH 7.4, 150 mM KCl, 10 percent glycerol, and 250 mM imidazole. Proteins were dialyzed overnight using 3 mL Slide-A-Lyzer 20K and 10K MWCO dialysis cassettes for MBP-FUS and MBP-TEV, respectively, (Thermo Scientific 66003 and 66455) with two rounds of 1 L dialysis buffer. For MBP-FUS, dialysis buffer consisted of 20 mM HEPES pH 7.4, 150 mM NaCl, 5 percent glycerol, and 5 mM beta-mercaptoethanol; for MBP-TEV, dialysis buffer consisted of 25 mM HEPES pH 7.4, 150 mM KCl, 10 percent glycerol, and 5 mM beta-mercaptoethanol.

Approximately 0.5 mL of purified MBP-FUS protein was fluorescently labeled with amine-reactive Atto 488-NHS ester (Sigma-Aldrich 41698-1MG-F). After resuspending the reactive dye in anhydrous DMSO to a stock concentration of 10 mM, dye was added to the protein at a dye:protein molar ratio of 1:1. The dye was allowed to conjugate to the protein for 20 minutes at room temperature, and the mixture was transferred to a 0.5 mL Slide-A-Lyzer 20K MWCO dialysis cassette (Thermo Scientific 66005) and dialyzed with two round of 0.5 L MBP-FUS dialysis buffer to remove unconjugated dye.

Protein and conjugated dye concentrations were measured using a Nanodrop spectrophotometer. The A260/A280 absorbance ratios were found to be in the range of 0.60-0.65, indicating relatively minor absorbance contributions from residual nucleic acids. Small aliquots of labeled and unlabeled proteins were snap frozen in liquid nitrogen and stored at -80°C.

### NSP13 protein production

#### Plasmid construction

The coding sequence for NSP13 from the SARS CoV-2 Washington isolate (Genbank MN985325) was synthesized as an E. coli codon-optimized fragment (GenScript) and cloned into the pUC57 vector by the manufacturer. The NSP13 sequence was PCR-amplified with Pfu polymerase, and the resulting fragment cloned into the BsaI site of the pSUMO (LifeSensors) plasmid using the HiFi Assembly kit (NEB) to produce an N-terminal six histidine-tagged SUMO- NSP13 fusion cassette (6XHis-SUMO-NSP13). The ULP-1 cleavage site encoded in the SUMO tag was located between the 6X-His-SUMO fragment and the NSP13 sequence to allow for the production of authentic N-terminal SARS Co-V2 NSP13 protein. Samples were sequenced at the UAMS Sequencing Core Facility using a 3130XL Genetic Analyzer (Applied Biosystems, Foster City, CA).

#### Sequence of NSP13

SUMO-NSP13 that is expressed, and the NSP13 WT in bold:

**MGHHHHHHGSDSEVNQEAKPEVKPEVKPETHINLKVSDGSSEIFFKIKKTTPLRRLMEAFAKRQ GKEMDSLRFLYDGIRIQADQAPEDLDMEDNDIIEAHREQIGG**AVGACVLCNSQTSLRCGACIRR PFLCCKCCYDHVISTSHKLVLSVNPYVCNAPGCDVTDVTQLYLGGMSYYCKSHKPPISFPLCANGQVFGLYKNTCVGSDNVTDFNAIATCDWTNAGDYILANTCTERLKLFAAETLKATEETFKLSYGIATVREVLSDRELHLSWEVGKPRPPLNRNYVFTGYRVTKNSKVQIGEYTFEKGDYGDAVVYRGTTTYKLNVGDYFVLTSHTVMPLSAPTLVPQEHYVRITGLYPTLNISDEFSSNVANYQKVGMQKYSTLQGPPGTGKSHFAIGLALYYPSARIVYTACSHAAVDALCEKALKYLPIDKCSRIIPARARVECFDKFKVNSTLEQYVFCTVNALPETTADIVVFDEISMATNYDLSVVNARLRAKHYVYIGDPAQLPAPRTLLTKGTLEPEYFNSVCRLMKTIGPDMFLGTCRRCPAEIVDTVSALVYDNKLKAHKDKSAQCFKMFYKGVITHDVSSAINRPQIGVVREFLTRNPAWRKAVFISPYNSQNAVASKILGLPTQTVDSSQGSEYDYVIFTQTTETAHSCNVNRFNVAITRAKVGILCIMSDRDLYDKLQFTSLEI PRRNVATLQ

#### Protein Expression and Purification

Wild-type plasmids were transformed into Rosetta2 cells, and colonies were grown to saturation overnight at 37°C in NZCYM (Research Products International) supplemented with kanamycin (50 µg/ml) and chloramphenicol (25 µg/ml). The cultures were diluted 1:100 in fresh antibiotic-containing NZCYM media and allowed to grow until reaching an OD600 nm of 0.8-1. The cultures were supplemented with 0.1 mM ZnSO4 and 0.2% dextrose and cooled on ice for 10 minutes. Expression of the proteins was induced with 0.2 mM isopropyl β-D-1- thiogalactopyranoside (IPTG) at 18°C for 12-16 hours. The cells were harvested by centrifugation at 4,000 x g for 15 minutes at 4°C, and the cell pellets were stored at -80°C until purification.

All purifications steps were carried out on ice or at 4°C. Pellets were fully resuspended in lysis buffer (50 mM sodium phosphate, pH 8.0, 300 mM NaCl, 1 mM β-mercaptoethanol, 10% glycerol and 20 mM imidazole) supplemented with 2 mM phenylmethylsulfonyl fluoride (PMSF) and 1X EDTA-free protease inhibitor cocktail (Pierce), lysed by microfluidization and clarified by centrifugation at 17,000 x g for 1 hour at 4°C. The His-tagged SUMO-NSP13 was initially isolated from the crude supernatant using immobilized metal ion affinity chromatography (IMAC). The sample was passed through a HisTrap FF column (Cytiva) at 1 ml/minute. The Ni Sepharose affinity resin was washed extensively with 20 column volumes of buffer, and the protein eluted with 10 column volumes of lysis buffer containing 200 mM imidazole. After dialysis of the pooled SUMO-NSP13-containing fractions into low imidazole lysis buffer, the 6XHis-SUMO tag was cleaved with 6XHis-ULP-1 for 4 hours at 4°C, and complete digestion was confirmed by SDS-PAGE analysis. The ULP-1 and SUMO tags were separated from the NSP13 proteins by subjecting the sample to a second round of Ni2+-affinity chromatography as before. NSP13- containing fractions were pooled, dialyzed against low salt buffer (50 mM sodium phosphate, pH 6.8, 150 mM NaCl, 4 mM β-mercaptoethanol, 0.5 mM EDTA and 10% glycerol) and passed through a HighTrap SP (Cytiva) ion exchange column. Under these conditions, NSP13 and mutants did not adhere to the SP column and the flow-thru fractions were collected. The NSP13- containing fractions were pooled and concentrated with an Amicon Ultra-15 centrifugation filter unit. The NSP13 protein sample was passed through a Sephacyl S200-HR HiPrep 26/60 (Cytiva) column equilibrated in NSP13 Storage Buffer (25 mM HEPES, pH 7.5, 150 mM NaCl, 0.5 mM TCEP and 20% glycerol). Purified NSP13 was quantified by UV spectrophotometry at 280 nm using the expected extinction coefficient of 68,785 M-1 cm-1 and confirmed using the BCA Protein Assay (Pierce). Protein samples were aliquoted and stored at -80°C.

##### RNA template production

RNAs representing the TRS-L (plus) and 6 of the TRS-Bs (plus and minus) of SARS-CoV-2 were produced by PCR amplification with T7 containing primers (iProof) from SARS-CoV-2 RT-PCR genome fragments 1, 4, and 5, a kind gift from Ahmet Yildez lab^22^. Templates were chosen to be equal length (480+ GGG) and have equal flanking sequences 5′ and 3′ of the TRS-B. Ensemble diversity was predicted using Vienna fold^95^. RNA–RNA interaction strength was predicted utilizing RNA-hybrid^45^. G-quadruplexes for both the plus and minus strand SARS-CoV-2 RNA genome were predicted using QGRS mapper^63^ under default parameters. The following template sequences were utilized, position of TRS sequence is marked in red and YRRRY’s is marked in yellow highlight.

**Spike plus** GGGGGTTATGTCATGCATGCAAATTACATATTTTGGAGGAATACAAATCCAATTCAGTTGTCTTCCTATTCTTTATTTGACATGAGTAAATTTCCCCTTAAATTAAGGGGTACTGCTGTTATGTCTTTAAAAGAAGGTCAAATCAATGATATGATTTTATCTCTTCTTAGTAAAGGTAGACTTATAATTAGAGAAAACAACAGAGTTGTTATTTCTAGTGATGTTCTTGTTAACAACTAAACGAACAATGTTTGTTTTTCTTGTTTTATTGCCACTAGTCTCTAGTCAGTGTGTTAATCTTACAACCAGAACTCAATTACCCCCTGCATACACTAATTCTTTCACACGTGGTGTTTATTACCCTGACAAAGTTTTCAGATCCTCAGTTTTACATTCAACTCAGGACTTGTTCTTACCTTTCTTTTCCAATGTTACTTGGTTCCATGCTATACATGTCTCTGGGACCAATGGTACTAAGAGGTT

**Envelope plus** GGGGTTGTATTACACAGTTACTTCACTTCAGACTATTACCAGCTGTACTCAACTCAATTGAGTACAGACACTGGTGTTGAACATGTTACCTTCTTCATCTACAATAAAATTGTTGATGAGCCTGAAGAACATGTCCAAATTCACACAATCGACGGTTCATCCGGAGTTGTTAATCCAGTAATGGAACCAATTTATGATGAACCGACGACGACTACTAGCGTGCCTTTGTAAGCACAAGCTGATGAGTACGAACTTATGTACTCATTCGTTTCGGAAGAGACAGGTACGTTAATAGTTAATAGCGTACTTCTTTTTCTTGCTTTCGTGGTATTCTTGCTAGTTACACTAGCCATCCTTACTGCGCTTCGATTGTGTGCGTACTGCTGCAATATTGTTAACGTGAGTCTTGTAAAACCTTCTTTTTACGTTTACTCTCGTGTTAAAAATCTGAATTCTTCTAGAGTTCCTGATCTTCTGGTCTAA

**Membrane plus** GGGTTATGTACTCATTCGTTTCGGAAGAGACAGGTACGTTAATAGTTAATAGCGTACTTCTTTTTCTTGCTTTCGTGGTATTCTTGCTAGTTACACTAGCCATCCTTACTGCGCTTCGATTGTGTGCGTACTGCTGCAATATTGTTAACGTGAGTCTTGTAAAACCTTCTTTTTACGTTTACTCTCGTGTTAAAAATCTGAATTCTTCTAGAGTTCCTGATCTTCTGGTCTAAACGAACTAAATATTATATTAGTTTTTCTGTTTGGAACTTTAATTTTAGCCATGGCAGATTCCAACGGTACTATTACCGTTGAAGAGCTTAAAAAGCTCCTTGAACAATGGAACCTAGTAATAGGTTTCCTATTCCTTACATG GATTTGTCTTCTACAATTTGCCTATGCCAACAGGAATAGGTTTTTGTATATAATTAAGTTAATTTTCCTCTGGCTGTTATGGCCAGTAACTTTAGCTTGTTTTGTGCT

**ORF7 plus** GGGATTCCAGTAGCAGTGACAATATTGCTTTGCTTGTACAGTAAGTGACAACAGATGTTTCATCTCGTTGACTTTCAGGTTACTATAGCAGAGATATTACTAATTATTATGAGGACTTTTAAAGTTTCCATTTGGAATCTTGATTACATCATAAACCTCATAATTAAAAATTTATCTAAGTCACTAACTGAGAATAAATATTCTCAATTAGATGAAGAGCAACCAATGGAGATTGATTAAACGAACATGAAAATTATTCTTTTCTTGGCACTGATAACACTCGCTACTTGTGAGCTTTATCACTACCAAGAGTGTGTTAGAGGTACAACAGTACTTTTAAAAGAACCTTGCTCTTCTGGAACATACGAGGGCAATTCACCATTTCATCCTCTAGCTGATAACAAATTTGCACTGACTTGCTTTAGCACTCAATTTGCTTTTGCTTGTCCTGACGGCGTAAAACACGTCTATCAGTTACGTGCC

**ORF8 plus** GGGGTTCATCAGACAAGAGGAAGTTCAAGAACTTTACTCTCCAATTTTTCTTATTGTTGCGGCAATAGTGTTTATAACACTTTGCTTCACACTCAAAAGAAAGACAGAATGATTGAACTTTCATTAATTGACTTCTATTTGTGCTTTTTAGCCTTTCTGCTATTCCTTGTTTTAATTATGCTTATTATCTTTTGGTTCTCACTTGAACTGCAAGATCATAATGAAACTTGTCACGCCTAAACGAACATGAAATTTCTTGTTTTCTTAGGAATCATCACAACTGTAGCTGCATTTCACCAAGAATGTAGTTTACAGTCATGTACTCAACATCAACCATATGTAGTTGATGACCCGTGTCCTATTCACTTCTATTCTAAATGGTATATTAGAGTAGGAGCTAGAAAATCAGCACCTTTAATTGAATTGTGCGTGGATGAGGCTGGTTCTAAATCACCCATTCAGTACATCGATATCGGTAATTAT

**Nucleocapsid plus** GGGAAATGGTATATTAGAGTAGGAGCTAGAAAATCAGCACCTTTAATTGAATTGTGCGTGGATGAGGCTGGTTCTAAATCACCCATTCAGTACATCGATATCGGTAATTATACAGTTTCCTGTTTACCTTTTACAATTAATTGCCAGGAACCTAAATTGGGTAGTCTTGTAGTGCGTTGTTCGTTCTATGAAGACTTTTTAGAGTATCATGACGTTCGTGTTGTTTTAGATTTCATCTAAACGAACAAACTAAAATGTCTGATAATGGACCCCAAAATCAGCGAAATGCACCCCGCATTACGTTTGGTGGACCCTCAGATTCAACTGGCAGTAACCAGAATGGAGAACGCAGTGGGGCGCGATCAAAACAACGTCGGCCCCAAGGTTTACCCAATAATACTGCGTCTTGGTTCACCGCTCTCACTCAACATGGCAAGGAAGACCTTAAATTCCCTCGAGGACAAGGCGTTCCAATTAACACCA

**Spike minus** GGGAACCTCTTAGTACCATTGGTCCCAGAGACATGTATAGCATGGAACCAAGTAACATTGGAAAAGAAAGGTAAGAACAAGTCCTGAGTTGAATGTAAAACTGAGGATCTGAAAACTTTGTCAGGGTAATAAACACCACGTGTGAAAGAATTAGTGTATGCAGGGGGTAATTGAGTTCTGGTTGTAAGATTAACACACTGACTAGAGACTAGTGGCAATAAAACAAGAAAAACAAACATTGTTCGTTTAGTTGTTAACAAGAACATCACTAGAAATAACAACTCTGTTGTTTTCTCTAATTATAAGTCTACCTTTACTAAGAAGAGATAAAATCATATCATTGATTTGACCTTCTTTTAAAGACATAACAGCAGTACCCCTTAATTTAAGGGGAAATTTACTCATGTCAAATAAAGAATAGGAAGACAACTGAATTGGATTTGTATTCCTCCAAAATATGTAATTTGCATGCATGACATAACC

**Envelope minus** GGGTTAGACCAGAAGATCAGGAACTCTAGAAGAATTCAGATTTTTAACACGAGAGTAAACGTAAAAAGAAGGTTTTACAAGACTCACGTTAACAATATTGCAGCAGTACGCACACAATCGAAGCGCAGTAAGGATGGCTAGTGTAACTAGCAAGAATACCACGAAAGCAAGAAAAAGAAGTACGCTATTAACTATTAACGTACCTGTCTCTTCCGAAACGAATGAGTACATAAGTTCGTACTCATCAGCTTGTGCTTACAAAGGCACGCTAGTAGTCGTCGTCGGTTCATCATAAATTGGTTCCATTACTGGATTAACAACTCCGGATGAACCGTCGATTGTGTGAATTTGGACATGTTCTTCAGGCTCATCAACAATTTTATTGTAGATGAAGAAGGTAACATGTTCAACACCAGTGTCTGTACTCAATTGAGTTGAGTACAGCTGGTAATAGTCTGAAGTGAAGTAACTGTGTAATACAAC

**Membrane minus** GGGAGCACAAAACAAGCTAAAGTTACTGGCCATAACAGCCAGAGGAAAATTAACTTAATTATATACAAAAACCTATTCCTGTTGGCATAGGCAAATTGTAGAAGACAAATCCATGTAAGGAATAGGAAACCTATTACTAGGTTCCATTGTTCAAGGAGCTTTTTAAGCTCTTCAACGGTAATAGT

ACCGTTGGAATCTGCCATGGCTAAAATTAAAGTTCCAAACAGAAAAACTAATATAATATTTAGTTCGTTTAGACCAGAAGATCAGGAACTCTAGAAGAATTCAGATTTTTAACACGAGAGTAAACGTAAAAAGAAGGTTTTACAAGACTCACGTTAACAATATTGCAGCAGTACGCACACAATCGAAGCGCAGTAAGGATGGCTAGTGTAACTAGCAAGAATACCACGAAAGCAAGAAAAAGAAGTACGCTATTAACTATTAACGTACCTGTCTCTTCCGAAACGAATGAGTACATAA

**ORF7 minus** GGGGGCACGTAACTGATAGACGTGTTTTACGCCGTCAGGACAAGCAAAAGCAAATTGAGTGCTAAAGCAAGTCAGTGCAAATTTGTTATCAGCTAGAGGATGAAATGGTGAATTGCCCTCGTATGTTCCAGAAGAGCAAGGTTCTTTTAAAAGTACTGTTGTACCTCTAACACACTCTTGGTAGTGATAAAGCTCACAAGTAGCGAGTGTTATCAGTGCCAAGAAAAGAATAATTTTCATGTTCGTTTAATCAATCTCCATTGGTTGCTCTTCATCTAATTGAGAATATTTATTCTCAGTTAGTGACTTAGATAAATTTTTAATTATGAGGTTTATGATGTAATCAAGATTCCAAATGGAAACTTTAAAAGTCCTCATAATAATTAGTAATATCTCTGCTATAGTAACCTGAAAGTCAACGAGATGAAACATCTGTTGTCACTTACTGTACAAGCAAAGCAATATTGTCACTGCTACTGGAAT

**ORF8 minus** GGGATAATTACCGATATCGATGTACTGAATGGGTGATTTAGAACCAGCCTCATCCACGCACAATTCAATTAAAGGTGCTGATTTTCTAGCTCCTACTCTAATATACCATTTAGAATAGAAGTGAATAGGACACGGGTCATCAACTACATATGGTTGATGTTGAGTACATGACTGTAAACTACATTCTTGGTGAAATGCAGCTACAGTTGTGATGATTCCTAAGAAAACAAGAAATTTCATGTTCGTTTAGGCGTGACAAGTTTCATTATGATCTTGCAGTTCAAGTGAGAACCAAAAGATAATAAGCATAATTAAAACAAGGAATAGCAGAAAGGCTAAAAAGCACAAATAGAAGTCAATTAATGAAAGTTCAATCATTCTGTCTTTCTTTTGAGTGTGAAGCAAAGTGTTATAAACACTATTGCCGCAACAATAAGAAAAATTGGAGAGTAAAGTTCTTGAACTTCCTCTTGTCTGATGAAC

**Nucleocapsid minus** GGGATAATTACCGATATCGATGTACTGAATGGGTGATTTAGAACCAGCCTCATCCACGCACAATTCAATTAAAGGTGCTGATTTTCTAGCTCCTACTCTAATATACCATTTAGAATAGAAGTGAATAGGACACGGGTCATCAACTACATATGGTTGATGTTGAGTACATGACTGTAAACTACATTCTTGGTGAAATGCAGCTACAGTTGTGATGATTCCTAAGAAAACAAGAAATTTCATGTTCGTTTAGGCGTGACAAGTTTCATTATGATCTTGCAGTTCAAGTGAGAACCAAAAGATAATAAGCATAATTAAAACAAGGAATAGCAGAAAGGCTAAAAAGCACAAATAGAAGTCAATTAATGAAAGTTCAATCATTCTGTCTTTCTTTTGAGTGTGAAGCAAAGTGTTATAAACACTATTGCCGCAACAATAAGAAAAATTGGAGAGTAAAGTTCTTGAACTTCCTCTTGTCTGATGAAC

**5′end plus** GGGTTAAAGGTTTATACCTTCCCAGGTAACAAACCAACCAACTTTCGATCTCTTGTAGATCTGTTCTCTAAACGAACTTTAAAATCTGTGTGGCTGTCACTCGGCTGCATGCTTAGTGCACTCACGCAGTATAATTAATAACTAATTACTGTCGTTGACAGGACACGAGTAACTCGTCTATCTTCTGCAGGCTGCTTACGGTTTCGTCCGTGTTGCAGCCGATCATCAGCACATCTAGGTTTCGTCCGGGTGTGACCGAAAGGTAAGATGGAGAGCCTTGTCCCTGGTTTCAACGAGAAAACACACGTCCAACTCAGTTTGCCTGTTTTACAGGTTCGCGACGTGCTCGTACGTGGCTTTGGAGACTCCGTGGAGGAGGTCTTATCAGAGGCACGTCAACATCTTAAAGATGGCACTTGTGGCTTAGTAGAAGTTGAAAAAGGCGTTTTGCCTCAACTTGAACAGCCCTATGTGTTCATCAAA

**Spike minus 10% Overlap** GGGTGGATTTGTATTCCTCCAAAATATGTAATTTGCATGCATGACATAACCATCTATTTGTTCGCGTGGTTTGCCAAGATAATTACATCCAATTAAAAATGCTTCAGATGATGACGCATTCACATTAGTAACAAAGGCTGTCCACCATGCGAAGTGTCCCATGAGCTTATAAAGATCAGCATTCCAAGAATGTTCTGTTATCTTTATAGCCACGGAACCTCCAAGAGCTAGCTTTTGTTGTATAAACCCACAAATGTAAGTGAAAAAACCCTCTTTAGAGTCATTTTCTTTTGTAACATTTTTAGTCTTAGGGTCGTACATATCACTAATAATGAGATCCCATTTATTAGCTGTATGTACAGTTGCACAATCACCAATCAAAGTTGAATCTGCATCAGAGACAAAGTCATTAAGATCTGAATCGACAAGCAGCGTACCCGTAGGCAACCACTGTCTTAAAACAGCTGTACCTGGTGCAACTCC

#### Nucleocapsid minus 10% Overlap

GGGTTCAATTAAAGGTGCTGATTTTCTAGCTCCTACTCTAATATACCATTTAGAATAGAAGTGAATAGGACACGGGTCATCAACTACATATGGTTGATGTTGAGTACATGACTGTAAACTACATTCTTGGTGAAATGCAGCTACAGTTGTGATGATTCCTAAGAAAACAAGAAATTTCATGTTCGTTTAGGCGTGACAAGTTTCATTATGATCTTGCAGTTCAAGTGAGAACCAAAAGATAATAAGCATAATTAAAACAAGGAATAGCAGAAAGGCTAAAAAGCACAAATAGAAGTCAATTAATGAAAGTTCAATCATTCTGTCTTTCTTTTGAGTGTGAAGCAAAGTGTTATAAACACTATTGCCGCAACAATAAGAAAAATTGGAGAGTAAAGTTCTTGAACTTCCTCTTGTCTGATGAACAGTTTAGGTGAAACTGATCTGGCACGTAACTGATAGACGTGTTTTACGCCGTCAGGACAA

### RNA PRODUCTION

RNA was produced according to our established protocols with T7 in vitro transcription kit (NEB) with labeled cy3, cy5, and atto488 UTPs and purified with LiCl precipitation. Produced RNAs were quality controlled for concentration (nanodrop) as well as size and purity on a denaturing 1% agarose gel (stained with sybr gold). Data for the manuscript are representative of at least 2 independent batches of RNA for each tested sequence (1 labeled and 1 unlabeled). Label fraction for RNA was quantified via nanodrop and RNAs. Prior to use, RNAs were diluted to 1μM stock solutions and stored in labeled aliquots in the -80°C.

#### DEPC RNA structure probing

To validate RNA-RNA interaction arising from RNA complementarity, we performed a series of chemical probing experiments coupled with Nanopore direct RNA sequencing. First, all the RNAs are prepared using in-vitro transcription as described above. Then, 250nM of each RNA is independently probed and probed with its counterpart (e.g. sense with antisense, anti- sense with 5′end) to discern how it samples its thermodynamic landscape independently and with a binding partner, respectively.

For the RNAs that are independently probed, 10μL of 250nM RNA was denatured at 90°C for 3 min, cooled on ice, and renatured in 150mM NaCl, 20mM HEPES ph 7.4 for 20 min at 20°C. To test RNA-RNA interaction, a 10μL stock containing 250nM of RNA1 and 250uM of its counterpart RNA will be denatured using the same process as described above. For (+) DEPC, RNA was treated with 10% DEPC for 45 min at 20°C. Diethyl pyrocarbonate (DEPC) (Sigma) carbethoxylates unpaired adenosine at N-6 or N-7 by opening the imidazole ring. The reactions were immediately purified using RNA Clean & Concentrator (Zymo Research) or RNA XP bead (Beckman Coulter).

Sequencing libraries were prepared following the direct RNA sequencing protocol RNA- 004 (Oxford Nanopore Technologies, ONT) using sequence specific 3′ adapters (which is made by annealing). After renaturation and treatment with DEPC, 3′ adapter-ligated RNAs were reverse-transcribed using superscript III. The reverse transcription step serves to prop up RNA using its complementary cDNA to obtain longer reads and minimize the impact of RNA structure on nanopore readthrough. (Note that the direct-RNA sequencing method is sequencing the RNA through Nanopore, not the cDNA) The reverse transcribed RNA was pooled for each experimental condition and a motor protein ligated, which were subjected to Oxford Nanopore sequencing using MinION Flow Cell (RNA) on GridION Mk1 for 72 hour duration. Reads were basecalled using Super accurate (SUP) model (Oxford Nanopore Technologies, ONT). Reads were aligned to target RNAs allowing for mismatches, insertions and deletions to account for nanopore sequencing errors as well as probing adducts.

For each read, we identify the mismatched nucleotides. Then, we calculated the reactivity for each adenosine by using the equation below:

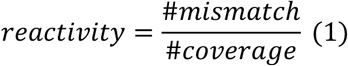

The uncertainty of reactivity is calculated using the standard error of binomial distribution, as the higher the read coverage associate with lower uncertainty:

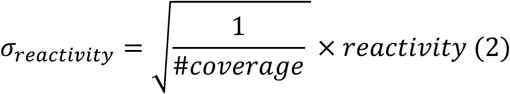

By comparing the DEPC (+) and DEPC (-) of the independently probed RNA, we concluded that the mismatch rate is a reliable way to gauge RNA structuredness, since the reactivity of the nucleotides on average is higher than the Nanopore sequencing error as reflected in DEPC (-). In addition, the ViennaRNA predicted centroid structure single stranded nucleotides has a higher reactivity on average than the double stranded nucleotides for all RNAs in question.

Next, we compared RNA when probed independently versus with its corresponding binding partner to investigate the nucleotides that are interacting with a binding partner, thereby protected from probing reagent.

For each adenosine position, the change in reactivity is calculated using:

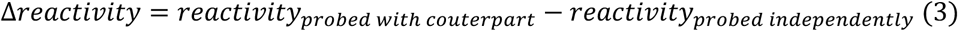

The uncertainty for the change in reactivity is calculated by propagating the uncertainty of the independent reactivity and uncertainty of reactivity when probed with partners, as described by the equation below

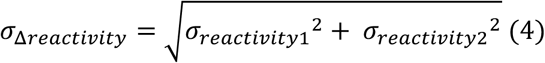

#### DEPC RNA structure models

RNA structure modeling was carried out with DEPC reactivity data and Viennafold software using the following parameters: Fold algorithms and basic options (MFE and partition function); Energy parameters (Turner model 2004); Energy parameters (37C and .150 M salt); After conversion of SHAPE reactivities (All loops Zarringhalam et al 2012); Convert SHAPE reactivities into pairing (Linear mapping).

### INOSINE CONTAINING RNA PRODUCTION

Inosine containing RNA was produced by modifying the in vitro transcription reaction replacing increasing percentages of ATPs with Inosine (cayman chemical).

### ERROR PRONE RNA PRODUCTION

Error prone PCR templates were produced (Jena) by utilizing multiple 30 cycle reactions with 1 μl of PCR product used as a template for the subsequent PCR reaction thereby increasing error rates. Only 4 to 5 cycles of error prone PCR resulted in difference in arrested network formation.

### CELL FREE CONDENSATION REACTION (N PROTEIN)

Condensates were formed in 384 well glass bottom dishes. 15 μl condensate buffer was added to the well (150mM NaCl, 20mM TRIS ph 7.4) followed by 5 μl of RNA and water (control water only). RNA, water, and condensate buffer were then mixed by pipetting a volume of 5μl three times. Following this, 5 μl of 16 μM (0.05% labeled) N protein in N protein buffer or N protein buffer only control (250 mM NaCl 20 mM phosphate buffer ph 7.4) was then added to the well and reactions were mixed by pipetting 5 μl 3 times. Reactions were incubated at 37^◦^C for 2-21 hours prior to imaging. For RNA combinations 1 μM stocks of labeled RNA were premixed on ice prior to addition unless staggering experiments were undertaken as in **Figure 4**.

For staggered RNA addition experiments, plus RNA was added followed by protein followed by minus after incubation. Cell free condensation reaction (FUS): FUS reactions were carried out in the same wells by precoating the plate with blocking solution 15 μl reaction buffer was added followed by 1 μl of TEV protease, followed by RNA followed by protein. A280 absorbance: diffuse phase measurements were acquired for each technical replicate by nanodrop with 1 μl of solution taken from the top of the well after at least 20 hours of incubation.

### CELL FREE CONDENSATION REACTION (N PROTEIN) WITH MG2+

Condensation was undertaking using staggered RNA addition with the addition of 10mM final concentration of MgCl2 to the reaction.

### IMAGE ACQUISITION

Images were acquired utilizing a Nikon spinning disc confocal with 100× silicone objective. Images are representative of at least 2 frames for at least 3 technical replicates. All images in each panel were acquired during the same imaging session with the same acquisition/laser settings. Displayed images in each panel are contrasted the same and the same area. Scale bars are indicated with a white line. Unless otherwise noted, images were acquired at the glass to be more comparable between round condensates which are found extend only a few microns in z to arrested networks which can extend 30+microns in z.

### IMAGE QUANTIFICATION

Condensate signal was thresholded on protein channel signal and 0.2μm pixel was used as a cutoff to detect particles. Region of Interest (ROI) maps from detected particles were used to extract size, shape (circularity), RNA and protein maximum intensity signal. Colocalization of RNA signals was quantified using colocalization threshold parameter for ImageJ.

### SIMULATION METHODS

For all simulations in this manuscript, we used LAMMPS^96^ molecular simulation package (version 23 Jun 2022). For all visualizations of simulation results, OVITO^97^ is used.

#### Coarse-grained Molecular Dynamics Simulations of Nucleocapsid protein and RNA (Model 1)

The nucleocapsid protein is modeled using the Mpipi model^40^. Since this model is mainly developed for studying the flexible disordered proteins, we used rigid bodies to describe the folded domains of N protein: RNA binding domain 1 (PDB:6YI3)^98^ and RNA binding domain 2 (PDB: 6YUN)^99^. Similarly, we have used full FUS protein (Uniprot ID: P35637) and treated RRM (281–369) and Zinc finger domain (452–443) using rigid bodies (with representative structures obtained from AlphaFold: AF-P35637-F1-v4). The protein–protein interactions are described by Wang–Frenkel potential (short-ranged non-bonded contacts) and Debye–Hückel theory (long-ranged electrostatics). While the original Mpipi force field contains parameters for RNA at the nucleotide resolution, the RNA is treated as fully flexible and are not able to form base pairs. On the other hand, the SIS RNA model^42^, which also represents RNA in a nucleotide level, supports description of RNA base-pairing^41,42^. The original SIS RNA model^42^ assumes that the electrostatic interactions are mainly screened out by divalent ions, leaving effective base-pair potentials to describe the interactions between nucleotides. More specifically, the base pairing is captured by a many-body potential that describes canonical A–U, G–C, and noncanonical G–U wobble pairing between nucleotides. The detailed form of the potential and the parameters are explicitly described in ^41^. Therefore, we have implemented a hybrid model that combines protein–protein and protein–RNA interactions from Mpipi with RNA–RNA interactions via a modified SIS RNA model. Specifically, we updated the RNA–RNA interactions with the SIS model parameters, and slightly adjusted the molecular diameters of nucleotide beads, such that they are consistent with the mapping used in the SIS RNA model. Moreover, since we are explicitly considering the electrostatics with Debye–Hückel screening, we updated the effective base pairing strengths under such consideration^100^.

In each simulation, N protein and the RNA chains are placed in cubic box with enough separation initially. Then each simulation is performed in the *NVT* ensemble using Langevin thermostat at 300 K for up to 2×10^8^ timesteps until the dimer formation is observed. The Debye screening length is set to 0.795 nm, which corresponds to a monovalent salt concentration of 0.15 M. The integration step size is set to 10 fs, and the friction is set to 𝛾= 0.01ps^-1^. Periodic boundary conditions are used.

For N protein–RNA contact maps, we monitor the inter-residue distance compared to the molecular diameters of nucleotides and amino acids, as set by the Wang–Frenkel 𝜎 parameter. Here, an amino acid 𝑖 is in contact with a nucleotide 𝑗, if distance 𝑟_&,7_ ≤ 1.2𝜎, which was empirically determined to be suitable for capturing key features in the contact matrices. For the N protein and RNA contact histogram, each distribution is normalized by the maximum number of contacts. For the NCPR (net charge per residue), a scanning window size of 5 amino acids are used.

To investigate the effect of excessive N protein or FUS protein on trans RNA contacts, we conduct two types of simulations for each protein. First, we perform simulations of 64 protein chains in the absence of RNA. Second, we simulate 64 protein chains with two complementary Envelope RNA strands. For both types of simulations, proteins and RNAs are premixed and placed closely in a cubic box of (600 Å)^3^. Then, the systems are simulated for up to 10^8^ timesteps using the exact same simulation parameters as described above for the N protein– RNA dimer simulations.

#### Coarse-grained Molecular Dynamics Simulations of RNA clusters (Model 2)

To investigate the relationship between RNA sequence and propensity to form trans RNA contacts, we simulate RNA strands in the absence of proteins. Here, we used the original SIS RNA model^41,42^, which was designed to capture the effect of RNA base pairing on RNA cluster formation at nucleotide resolution. Using this model, we study RNA-only systems, where the computational expense remains high even at nucleotide resolution due to the length of the RNA strands. Notably, probing the systems in the absence of proteins helps us to interrogate whether RNA–RNA interactions are sufficient to lead to the arrested phenotypes observed in our experiments. We simulated three scenarios for each sequence, a system with only 54 plus strands, a system with only 54 minus strands and a system with half plus (27) and half minus (27) strands. In all simulations, overall concentration, annealing protocol, thermodynamic ensemble (*NVT*), and the damping coefficients are identical^41^. For simulating the Envelope RNA sequence with 162 chains, we use the same overall concentration for 1:1 and 5:1 plus/minus systems. We also characterized finite size effects for our simulations and concluded that 54- chain simulations are sufficient for capturing statistically robust results (**Supplementary Figure S3**).

Each simulation is performed for up to 1.2 × 10^8^ timesteps, where the integration step size is set to 10 fs, and the friction is set to *𝛾* = 0.01ps ^−1^ for the Langevin thermostat at 293 K (following ^42^). This yields a total simulation time of over 1us. Periodic boundary conditions are used, and a short annealing process is performed at the beginning of the simulations to relax the system before equilibration. The data from the last 100 ns is used to calculate the near/at equilibrium RNA–RNA interaction strength, number of cis and trans RNA contacts, and the mean-square-displacements of RNA strands in each system.

For obtaining the average interaction strength, we have obtained the potential energies 𝑈 from the trajectories, which are then normalized by the total number of nucleotides in the simulations. To determine the number of trans RNA contacts, we evaluated the base-pair energies between nucleotides on different strands and define a trans RNA contact if the base-pairing energy is lower than −1.5𝐾_8_𝑇, which indicates the stability of the established contact against the thermal fluctuations. Finally, for the contact map analysis, we define contacts using the method described above (but for both intra- and inter-strand cases) and normalized the contact matrices by the maximum contacts of the corresponding simulations.

#### Limitations of wet-lab experiments

This study addresses the regulation of biomolecular condensation using purified minimal components of the SARS-CoV-2 virus. Many additional cellular and viral factors likely play important roles in this process.

#### Limitations of simulations

Coarse-grained simulation of protein and RNA is powerful as it can provide molecular insights into the studied system, while enabling us to probe longer timescales and larger system sizes, compared to all-atom simulations. However, these approximations may lead to certain limitations.

1. In all simulations, each amino acid or nucleotide is represented using a single bead or a single interaction site, which means that the structural and dynamic changes below this resolution are inaccessible from our simulations.
2. In representing the full N protein structure, we used the rigid-body representation for the RNA binding domains, under the assumption that such structured regions may experience minimal conformational changes under the simulation conditions. As a result, the simulation is not able to capture the effects that are related to structural changes in these domains.
3. In the study of N protein and RNA dimeric systems, we have found a nonspecific binding pattern that originates from the structure and the charge distribution of N protein. However, this model is not able to capture more specific binding interactions between N protein and RBD2, which may be important. Here, we capture mainly nonspecific binding effects.
4. Counterion, which is important to RNA condensation, is considered differently when proteins are involved (Model 1) and when RNA clustering is studied (Model 2). In the simulations when proteins are considered, the effective charges are explicitly modeled using Debye–Hückel theory as Mpipi force field is mainly parameterized under 0.15M NaCl concentrations. Therefore, the interaction sites in the RNA representation are also charged. However, when the RNA clustering is studied, it is assumed that the added monovalent and divalent ions effectively screen the electrostatic repulsions between the phosphate groups, and thus the remaining main interaction is the base-pairing between nucleotides at high salt concentrations.
5. In all simulations, only the canonical cis Watson-Crick (cWW) base pairing (GC and AU) and GU wobble pairs are considered. These interactions are favored in the model since cWW base pairing represents *>*70% of different pairing types in PDB structures and GC, AU, and GU pairing are the most dominant base pairs in cWW base pairing (*>*90%). However, effects of other base pairs could also serve to augment the degree of trans RNA contacts in our systems.

All relevant supporting data and codes are available in the Figshare data repository at: https://doi.org/10.6084/m9.figshare.28087196.

## SUPPLEMENTARY MATERIAL

**Supplemental Figure S1.**
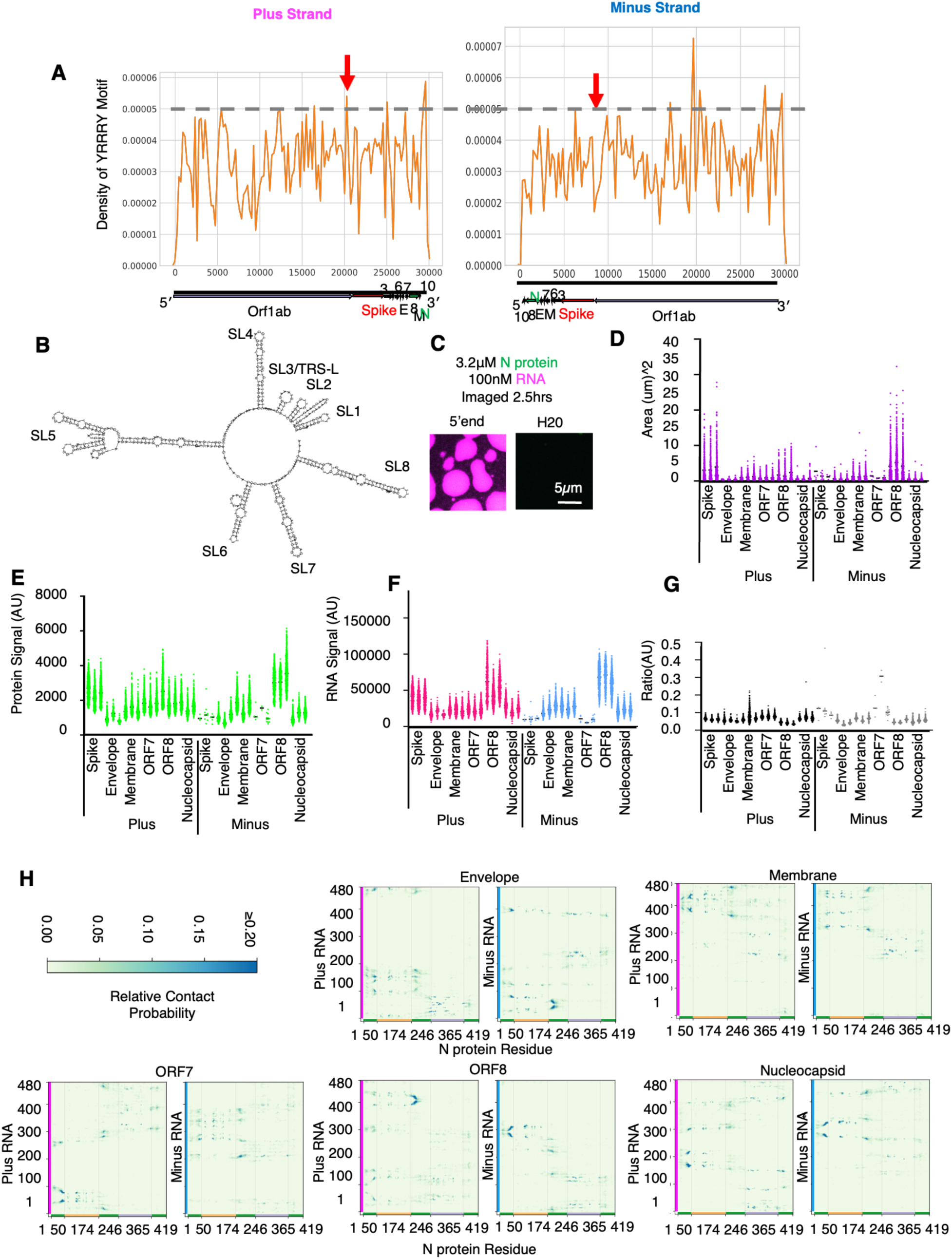
**TRS-N protein in reconstituted and simulated condensates.** Related to **Figure 1**. **(A)** The local density of YRRRY motifs on plus and minus strands. (**B**) Structure model for 5′end RNA (**C**) 5′end RNA condensates under tested conditions for TRS-B fragments from **Figure 1F**. **(D)** Condensate area (𝜇m2) quantification for **Figure 1F** taken from the protein signal. Data show the quantification of one representative image from 3 technical replicates. Plus Spike and minus ORF8 produce consistently larger condensates. Black line is the mean. **(E)** Measurement of max intensity of the protein signal for **Figure 1F**. **(F)** Measurement of max intensity of the RNA signal for panel **Figure 1F**. **(G)** Measurement of ratio max intensity of the RNA and protein signals for **Figure 1F**. **(H)** N protein and protein–RNA interactions are modeled via model 1 (see Methods). For each RNA, plus and minus strands are simulated with N protein separately. The corresponding domain boundaries of N protein are shown by vertical gray lines. The edges of the contact map are colored by the type of biomolecule/domain: yellow indicates RBD 1 of N protein, light purple for RBD 2 of N protein, green for disordered regions of N protein, purple for plus RNA strands, and blue for minus RNA strands. On the contact maps, the high probability region is indicated by bright blue colors. In terms of RNA sequences, the interaction hot spots appear to be random. However, as shown in main text **Figure 1**, the binding spots are relatively conserved with respect to the N protein, which is highly correlated with the charge distributions of N protein.

**Supplemental Figure S2.**
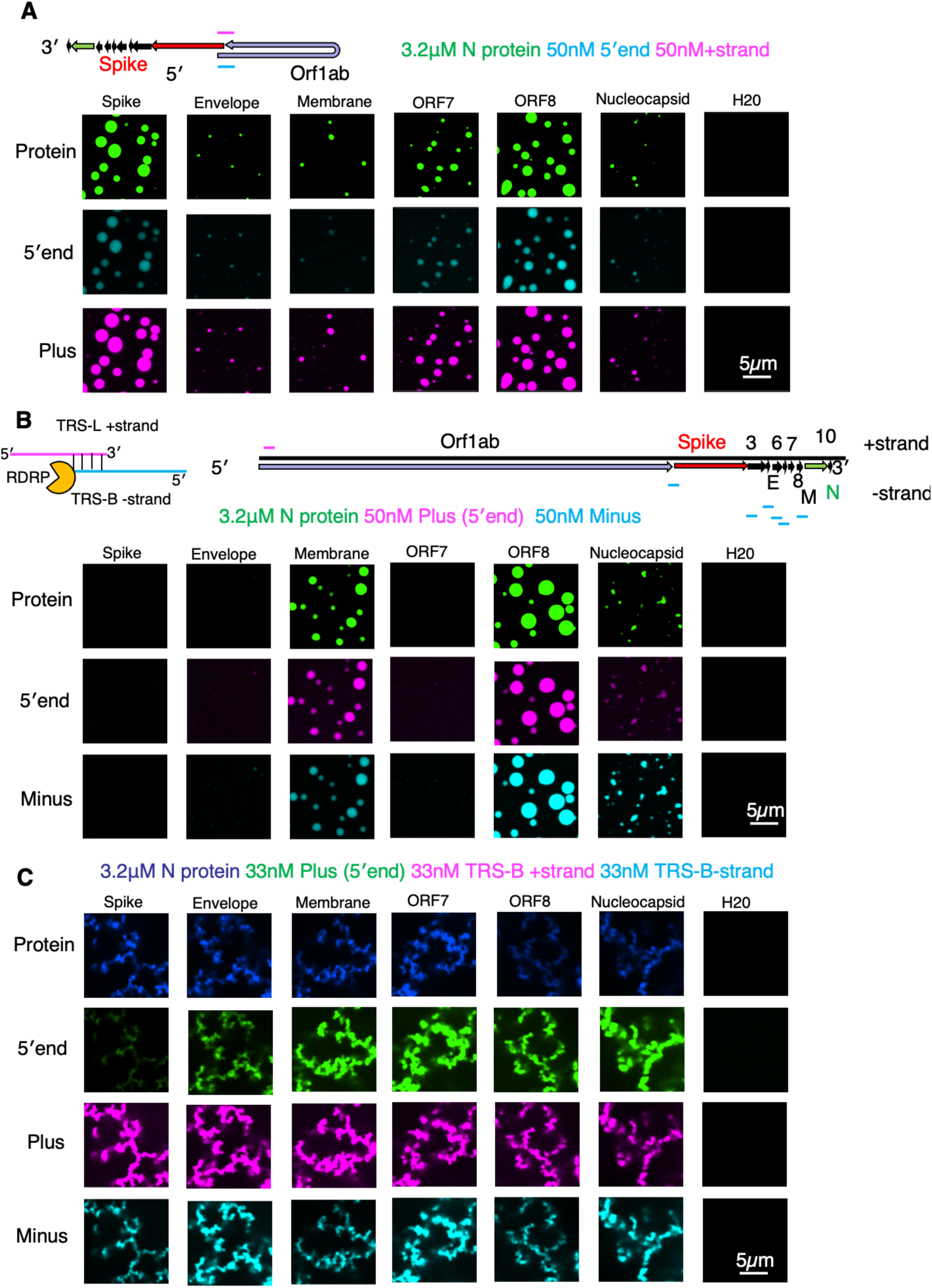
Separated RNA and protein channels from Figure 2. For **A-C**, All images depict the unmerged images related to **Figure 2** to indicate that all reconstituted RNAs and proteins are well mixed and present in the same condensate. Of note, **panel C**, N protein is labeled in blue rather than green and 5′end is instead labeled in green.

**Supplemental Figure S3.**
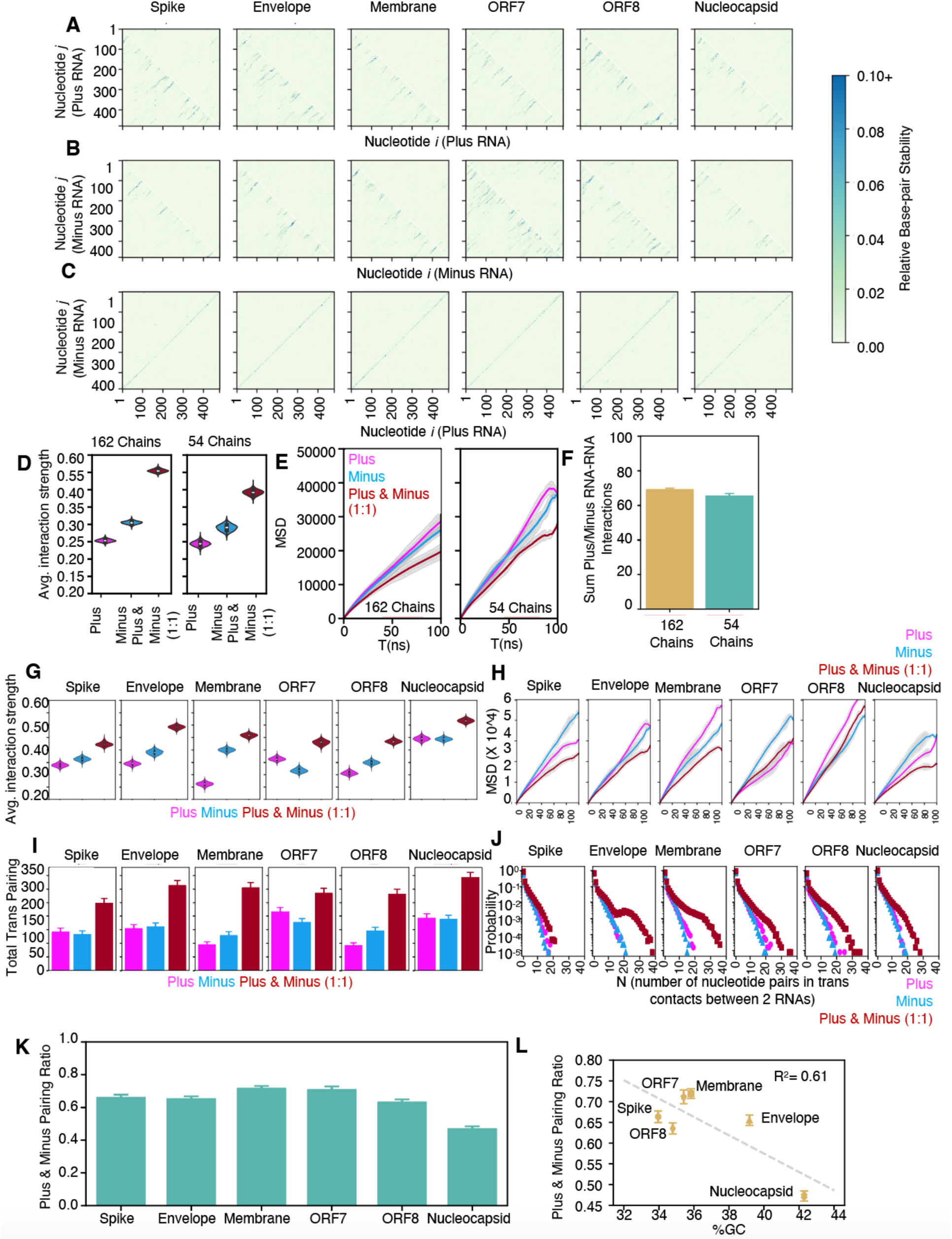
Simulations of RNA–RNA contacts, interaction strengths, and dynamics of RNA chains. Related to Figure 3. (**A**) Relative base pairing probability of intra-chain contacts (lower triangle) vs the inter-chain contacts (upper triangle) in plus RNA simulations. (**B**) Relative base pairing probability of intra-chain contacts (lower triangle) vs the inter-chain contacts (upper triangle) in minus RNA simulations. (**C**) Relative base-pair probability of inter-chain contacts in plus and minus 1:1 simulations. As expected, the contacts are along the diagonal due perfect complementary nature. (**D**) Comparison of the normalized interaction strength for different simulation sizes. The simulations are performed at the same concentration with periodic boundary conditions. The equilibrium potential energy absolute value |U| is normalized by calculating the average potential energy per nucleotide, |U|/N, where N is the number of nucleotides in the system (162x480 or 54x480). There is slight deviation of the values but the overall ordering of plus, minus, and 1:1 plus/minus systems are consistent between different system sizes. (**E**) Comparison of mean-squared displacement (MSD) of RNA chains for different systems sizes. The absolute values vary due to the number chains with 54-chain system moving faster for the same strands. However, the overall ordering is consistent. The pure plus and minus simulations result in an almost indistinguishable diffusion profile, and in both system sizes the movement of chains in the 1:1 plus/minus mixture system are noticeably reduced. (**F**) The percentage of plus– minus interactions in simulations of different sizes. This is obtained by dividing the plus–minus inter-chain base-pairing energies by overall inter-chain base-pairing energies. In both cases, the plus–minus pairing accounts for more than 60% inter-chain base-pairing interactions. (**G**) Average RNA–RNA interaction strength. This is obtained by normalizing the potential energy of each system by the total number of nucleotides. (**H**) Comparison of the MSD of the center of mass of chains. Here, the 1:1 plus/minus mixture exhibits the slowest dynamics for all strands. (**I**) Average number of nucleotides pairs in trans RNA contacts. (**J**) Probability distribution of number of nucleotide pairs in trans RNA contacts. (**K**) Comparison of base pairing ratio between simulations of 1:1 plus/minus systems. These values are obtained by dividing the plus–minus inter-chain base-pairing energies by overall inter-chain base-pairing energies. This ratio is over 50% for all the cases, except for nucleocapsid RNAs. (**L**) The percentage of GC in RNA sequences and base pairing ratio between simulations of 1:1 plus/minus mixtures. These two quantities are overall negatively correlated, indicating that GC pairing plays an important role in 1:1 plus/minus mixture inter-chain pairing.

**Supplemental Figure S4.**
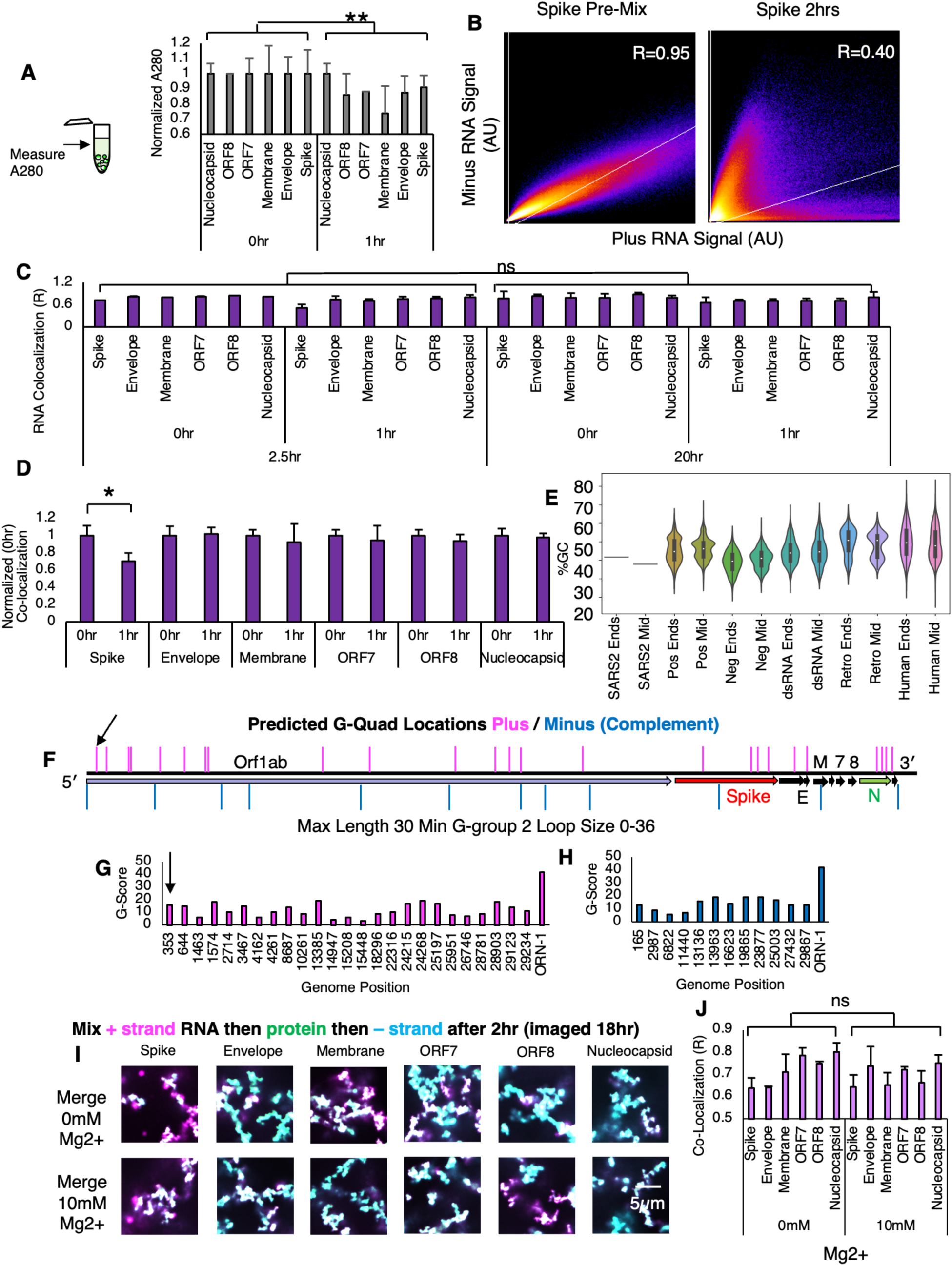
Mixing of time-delayed RNA additions is controlled by GC content. Related to **Figure 4**.**(A)** A280, diffuse phase measurements of RNA and protein signal taken from **Figure 4C**. Consistent with plus TRS-B containing fragments being better able to drive N protein condensation (**Figure 1**, top panel), Reactions with longer incubation with plus RNA only (1 hour) have lower diffuse phase signal than 0-hour conditions (p*<*0.01 **, P*<*0.05 *). **(B)** Representative plots (Spike) taken from the colocalization calculation. For the plus (x-axis) and minus (y-axis) RNA signals for RNA premixed on ice (left panel) and minus RNA mixed 2 hours post plus and N protein mixing. Premixed RNAs show a higher degree of co-localization with R values approaching 1. Lack of co-localization is observed by staggering addition and low R values. **(C)** Related to **Figure 4B** and **4C**. Minor changes for most RNA combinations following 18 hours of further incubation with minus RNA (images taken at 2.5 and 20 hours). No significant change in recorded values. **(D)** Related to **Figure 4C**. Normalized (0 hour) co- localization signal at 2.5 hours. The mixing of the highest GC RNA combinations is not enhanced by additional preincubation of N protein with plus RNA apart from spike. **(E)** Percent GC content in ends (200nt not including polyA) versus middle of the virus. SARS-CoV-2 shows higher GC content at ends on average with most RNA viruses and human transcripts showing the opposite enrichment. **(F)** G-quadruplexes were predicted using QGRS mapper using default conditions (Max length 30, Min G group 2, Loop size 0-36). Of note, these settings represent the least stringent possible criteria for g-quadruplex prediction. Pink lines indicate the approximate location of predicted G-quadruplexes on the plus stand and turquoise indicates the minus strand (in the complement configuration for easy comparison of duplex position sequences). Black arrow indicates the only predicted g-quadruplex which overlaps with a tested RNA fragment. 0/12 TRS-B containing fragments in both the plus and minus strand sequence had predicted g-quadruplexes. **(G)** Quality score for plus and minus strand RNAs of the SARS-CoV-2 genome. Each bar is labeled with the start position of the indicated g-quadruplex in the plus or minus **(H)** strand genome. Both plus and minus g-quadruplexes have much lower quality scores than bona fide g-quadruplex from Orn-1. **(I)** Addition of non-physiologically high levels of magnesium, which should support g-quadruplex formation, has no obvious impact on plus (magenta) and minus (cyan) strand RNA co-localization following staggered addition pre-incubation experiment with N protein (green). Plus strand RNA mixed with N protein followed by 2 hours of incubation followed by minus strand addition. **(J)** Quantification of the co-localization of the plus and minus strand RNA co-localization with or without additional Mg2+ indicating that there is no significant difference (ns). Error bars represent technical replicates.

**Figure S5.**
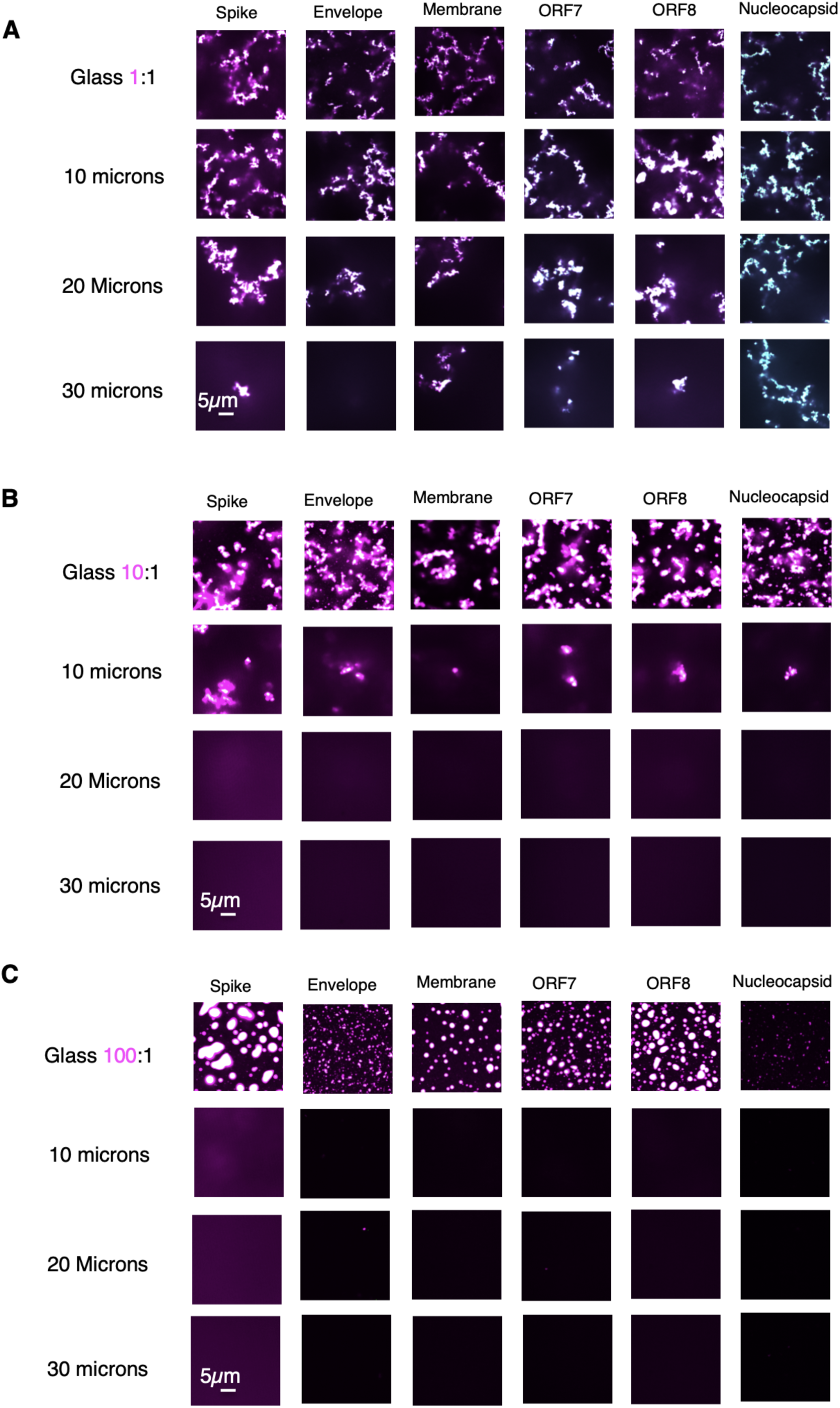
Arrested networks extend farther into Z than non-arrested networks. Related to Figure 5 **(A)** Representative merged images from 1:1 plus/minus RNA condition at glass (row 1), 10 (row 2), 20 (row 3), and 30 (row 4) microns in Z. All but envelope extends 30 microns in Z in these conditions. **(B)** Representative merged images from 10:1 plus/minus RNA condition at glass (row 1), 10 (row 2), 20 (row 3), and 30 (row 4) microns in Z. All extend 10 microns in Z in these conditions. **(C)** Representative merged images from 100:1 plus/minus RNA condition at glass (row 1), 10 (row 2), 20 (row 3), and 30 (row 4) microns in Z. No tested conditions extend 10 microns in Z in these conditions.

